# A Saccharimonadia epibiont interacts with the polyphosphate-accumulating bacterium *Candidatus* Phosphoribacter in wastewater treatment plants

**DOI:** 10.64898/2026.09.16.751625

**Authors:** Huifeng Hu, Katharina Kitzinger, Marta Nierychlo, Lei Liu, Anna Lopatina, Jannie Munk Kristensen, Miriam Peces, Craig William Herbold, Morten Kam Dahl Dueholm, Daan Speth, Holger Daims, Petra Pjevac, Per Halkjaer Nielsen, Michael Wagner

**Author notes:** Corresponding author: Michael Wagner.

## Abstract

Many Patescibacteria lineages are abundant in wastewater treatment plants, yet their interaction partners and ecological roles remain poorly understood. Here, we identify a specific association between a novel Saccharimonadia genus, *Candidatus* Lutisaccharimonas, and the widespread polyphosphate-accumulating organism (PAO) *Candidatus* Phosphoribacter. Fluorescence *in situ* hybridization revealed close physical proximity between the two taxa, supporting an epibiotic relationship. Correlated population dynamics, CRISPR spacer matches and evidence of horizontal gene transfer further corroborated this association. Comparative genomic analyses showed that *Ca.* Lutisaccharimonas has greater metabolic versatility than is typical of Patescibacteria, including complete pathways for *de novo* nucleotide biosynthesis and genes involved in amino acid metabolism. These metabolic features may shape its interactions with *Ca.* Phosphoribacter. Together, our findings identify a previously undescribed host association for Patescibacteria, expand current understanding of their metabolic potential, and suggest that the interactions between Saccharimonadia and PAOs may influence phosphorus removal in engineered ecosystems.

## Introduction

Patescibacteria are a monophyletic bacterial group characterized by ultrasmall cell sizes, reduced genomes, limited biosynthetic capabilities and highly divergent ribosomal RNA genes^1,2^. Saccharimonadia (previously referred to as TM7) is currently the most studied class of Patescibacteria and is detected across human^3^, animal^4^, and environmental^5^ samples as well as engineered systems^6^. Saccharimonadia often live as episymbionts of bacterial hosts, with interactions ranging from parasitism to mutualism^3,7–9^. In 2015, the first epibiotic Saccharimonadia species, *Nanosynbacter lyticus* (TM7x), was isolated from the human oral cavity together with its host *Actinomyces odontolyticus*^3^. *Actinomyces odontolyticus* and *Nanosynbacter lyticus* can coexist in a stable long-term association^10,11^, and the epibiont can even protect its host against acid stress in the human oral cavity, and from infection by lytic phages^8,12^. In activated sludge, the Saccharimonadia representative *Ca.* Mycosynbacter amalyticus has been shown to lyse foam-forming bacteria, such as *Gordonia amarae* and other mycobacterial species^7^, thus potentially regulating microbial community composition and affecting sludge bulking and foaming behavior.

In wastewater treatment plants (WWTPs), microbial consortia are widely employed for nutrient removal and resource recovery. The enhanced biological phosphorus removal (EBPR) process uses polyphosphate-accumulating organisms (PAOs) to remove phosphorus from wastewater^13^. It is based on the sequential and repeated exposure of activated sludge to anoxic and oxic conditions and leverages the distinctive metabolic traits of PAOs. In the anoxic phase, PAOs use polyphosphate (polyP)-derived energy to accumulate carbon-rich storage compounds like polyhydroxyalkanoates (PHA) and glycogen, and subsequently exploit these compounds as carbon and energy sources for growth and accumulation of polyP during the oxic phase. After every oxic phase, most of the activated sludge is pumped back into the anoxic basin, while excess biomass, including the stored polyP, is removed from the treated wastewater. Due to their biotechnological importance, PAOs have been extensively studied, and *Ca.* Accumulibacter has traditionally been regarded as the main PAO in WWTPs^14^. However, recent findings suggest that the genus *Ca.* Phosphoribacter also plays a significant functional role as a PAO in many EBPR systems^13,15,16^.

Our previous research on the global distribution patterns of Patescibacteria in WWTPs suggested, based on co-occurrence analyses, *Ca.* Phosphoribacter as a potential host organism for Saccharimonadia-related Patescibacteria^6^. In this study, we substantiated this hypothesis using fluorescence *in situ* hybridization (FISH) visualization and spatial coaggregation analysis in activated sludge samples from multiple WWTPs. We confirmed the epibiotic relationship of *Ca.* Phosphoribacter and members of an understudied Saccharimonadia lineage. We used the genome-based GTDB taxonomy for defining this lineage, which corresponds to genus GCA-2746885 within the family UBA4665 in GTDB release R220^17^. This genus has no cultured representatives and comprises organisms detected primarily in environmental and engineered ecosystems. We propose the genus name *Ca.* Lutisaccharimonas gen. nov. (Lu.ti.sac.cha.ri.mo’nas. L. n. _lutum_, dirt; N.L. n. _sacchara_, sugar; L. fem. n. _monas_, a unit, monad; N.L. fem. n. _Lutisaccharimonas_, a monad from dirt using sugar) for members of this clade and use this genus name throughout this study. In the 16S rRNA gene-based MiDAS taxonomy (Microbial Database for Activated Sludge)^15^, the 16S rRNA genes of members of this GTDB lineage were assigned to several closely related genera, but most of the sequences belong to the MiDAS placeholder genera midas_g_67 and midas_g_363. Thus, the boundaries of the genus are defined according to GTDB rather than the MiDAS assignments.

In addition to direct visualization, we examined correlations in the long-term population dynamics of *Ca.* Lutisaccharimonas and *Ca.* Phosphoribacter using weekly or biweekly time-series samples from three Danish EBPR plants. We further identified CRISPR spacer matches and evidence of horizontal gene transfer between the two taxa. Finally, comparative genomic analyses showed that *Ca.* Lutisaccharimonas genomes encode greater metabolic versatility than those of most other Patescibacteria, suggesting reduced host dependence and a potential capacity to exploit metabolites associated with carbon storage in *Ca*. Phosphoribacter.

## Results

### *In situ* detection of *Ca.* Lutisaccharimonas and coaggregation with *Ca.* Phosphoribacter in activated sludge samples

To determine whether *Ca.* Lutisaccharimonas and *Ca*. Phosphoribacter were physically associated in activated sludge, we designed two 16S rRNA-targeted oligonucleotide probes for *Ca.* Lutisaccharimonas (Sac732 and Sac1343, Supplementary Table 1) and applied them in FISH experiments with samples from four Danish full-scale EBPR WWTPs. Because full-length 16S rRNA gene sequences assigned to midas_g_67 and midas_g_363 were interspersed in phylogenetic analyses with sequences from closely related genera within the family midas_f_67, neither probe was completely specific for midas_g_67 and midas_g_363. However, the additionally targeted taxa occurred at very low relative abundances in the analyzed plants^18^, and overlapping signals from the two probes labeled with different fluorochromes confirmed the identity and morphology of the detected *Ca.* Lutisaccharimonas cells (Fig. 1A, Supplementary Figs. 1 and 2). Activated-sludge samples from WWTPs located in Ejby Mølle, Hjørring, Horsens and Aars were selected based on the abundance of 16S rRNA genes of *Ca.* Lutisaccharimonas and *Ca*. Phosphoribacter (Supplementary Fig. 2). Ethanol-fixed and hybridized *Ca.* Lutisaccharimonas cells were round-shaped, with mean dimensions of 0.8 × 0.5 μm. This size is considerably larger than that of *Ca*. Mycosynbacter amalyticus isolated from a WWTP or other isolated Saccharimonadia from the human oral cavity, but similar to that of environmental Saccharimonadia in freshwater^5^.

**Fig. 1.**
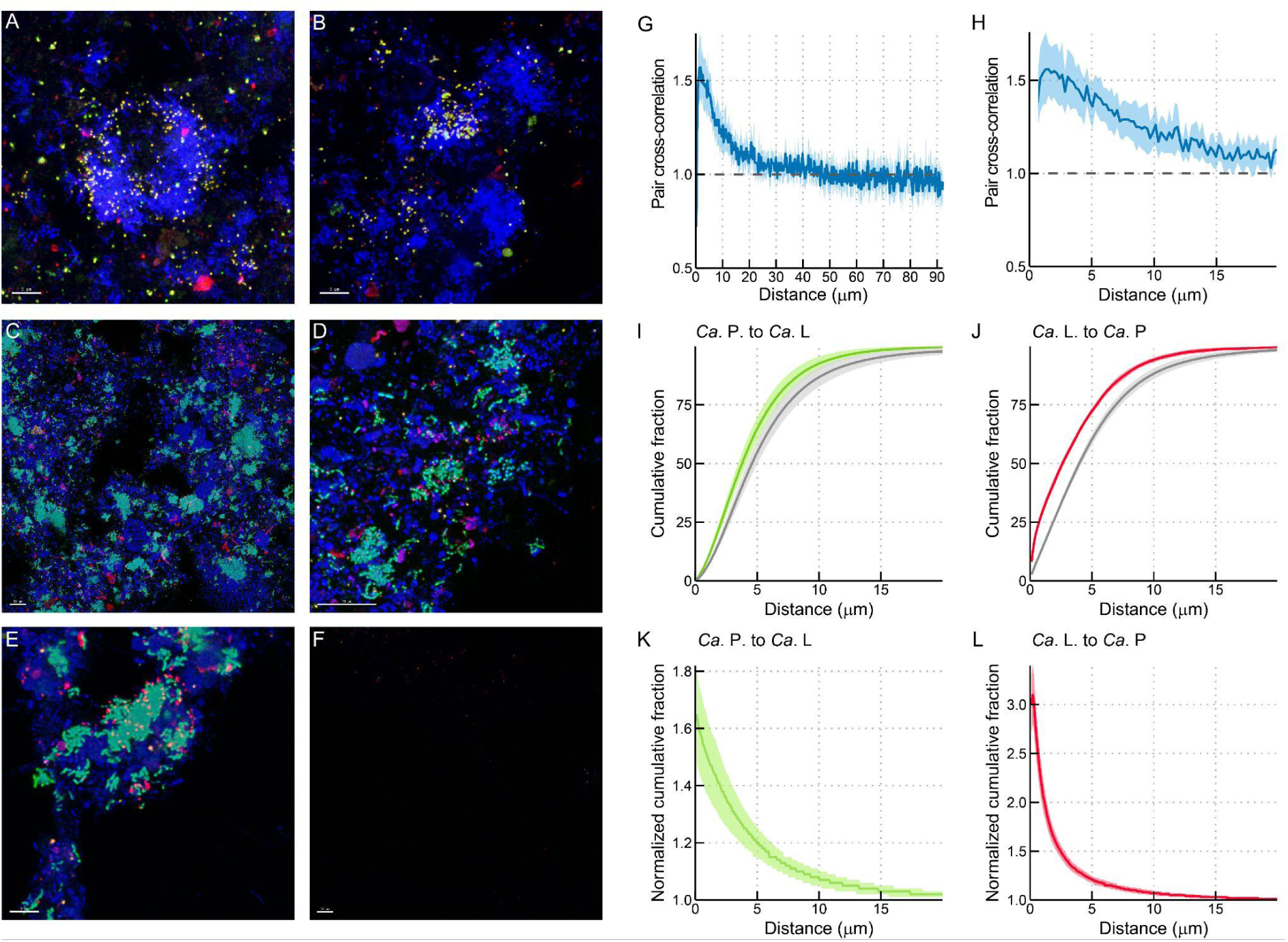
*Ca*. Lutisaccharimonas FISH visualization and coaggregation with *Ca.* Phosphoribacter. (A,B) Double hybridization of *Ca*. Lutisaccharimonas with probes Sac732 (Atto488/green) and Sac1343 (Cy3/red) in a sample from WWTP Horsens (A) and Hjørring (B). The *Ca.* Phosphoribacter-targeting probe Tet183 was used as a Cy5-labeled derivative (blue). Double-labeled *Ca*. Lutisaccharimonas cells appear yellow, *Ca.* Phosphoribacter cells appear blue. (C–E) Spatial coaggregation visualization of *Ca*. Lutisaccharimonas and *Ca.* Phosphoribacter*. Ca.* Phosphoribacter is labeled by probe Phos601 (Atto488/green) and *Ca*. Lutisaccharimonas is labeled by Sac1343 (Cy3/red). The EUB338 probe mix (Cy5) is shown in blue while *Ca.* Phosphoribacter and *Ca*. Lutisaccharimonas appear in cyan and magenta, respectively. Samples in C and D are from WWTP at Horsens, the sample in panel E is from WWTP at Ejby Mølle. (F) Negative control using nonEUB in Atto488 (green), Cy3 (red) and Cy5 (blue) in WWTP Horsens sample. The scale bars in panels A–F are 5 µm, 5 µm, 10 µm, 10 µm, 5 µm and 10 µm, respectively. (G,H) Spatial association between *Ca*. Lutisaccharimonas and *Ca.* Phosphoribacter in WWTP Horsens assessed by linear dipole analysis. The pair cross-correlation function *g*(*r*) > 1, *g*(*r*) = 1, *g*(*r*) <1 indicates coaggregation, random distribution, and segregation, respectively (see Methods for a detailed explanation). The solid blue line shows the mean of *g*(*r*), and the blue shading indicates 95% confidence limits. Panel G represents distances from 0 to 90 µm, and panel H represents distances from 0 to 20 µm. (I–L) Spatial association between *Ca*. Lutisaccharimonas (indicated by *Ca*. L in the figure) and *Ca.* Phosphoribacter (indicated by *Ca*. P in the figure) assessed by applying the inflate algorithm in both directions. In analyses which treated *Ca.* Phosphoribacter as the analyzed population and *Ca*. Lutisaccharimonas as the reference population, green shading indicates 95% confidence limits (I,K), while in the reciprocal analyses (J,L), red shading indicates 95% confidence limits. Panel I and panel J show the observed cumulative fractions of the analyzed population within increasing distances from the surface of the reference population. Colored curves represent the observed microbial populations, whereas gray curves (shading indicates 95% confidence limits) represent virtually simulated, randomly distributed populations of equivalent abundance. Panels K and L show normalized cumulative fractions, calculated as the cumulative fraction ratio of the observed to the virtual random populations. A normalized value of 1 represents spatial randomness, whereas values above 1 indicate coaggregation.

After validating the *Ca.* Lutisaccharimonas-targeted FISH probes, we combined probe Sac1343 with a modified Phos601 probe targeting both *Ca*. Phosphoribacter hodrii (Ca. P. hodrii) and *Ca*. Phosphoribacter baldrii (*Ca*. P. baldrii)^13^. *Ca.* Lutisaccharimonas cells in very close proximity to *Ca.* Phosphoribacter were detected in samples from all four WWTPs (Fig. 1C–E, Extended Data Fig. 1), indicating that the association is reproducible across distinct treatment plants. Negative controls using nonsense probes labeled with the same dyes were used for all four WWTPs, and in these experiments only background or small autofluorescence particles were observed (Fig. 1F and Extended Data Fig. 1).

**Extended Data Fig. 1.**
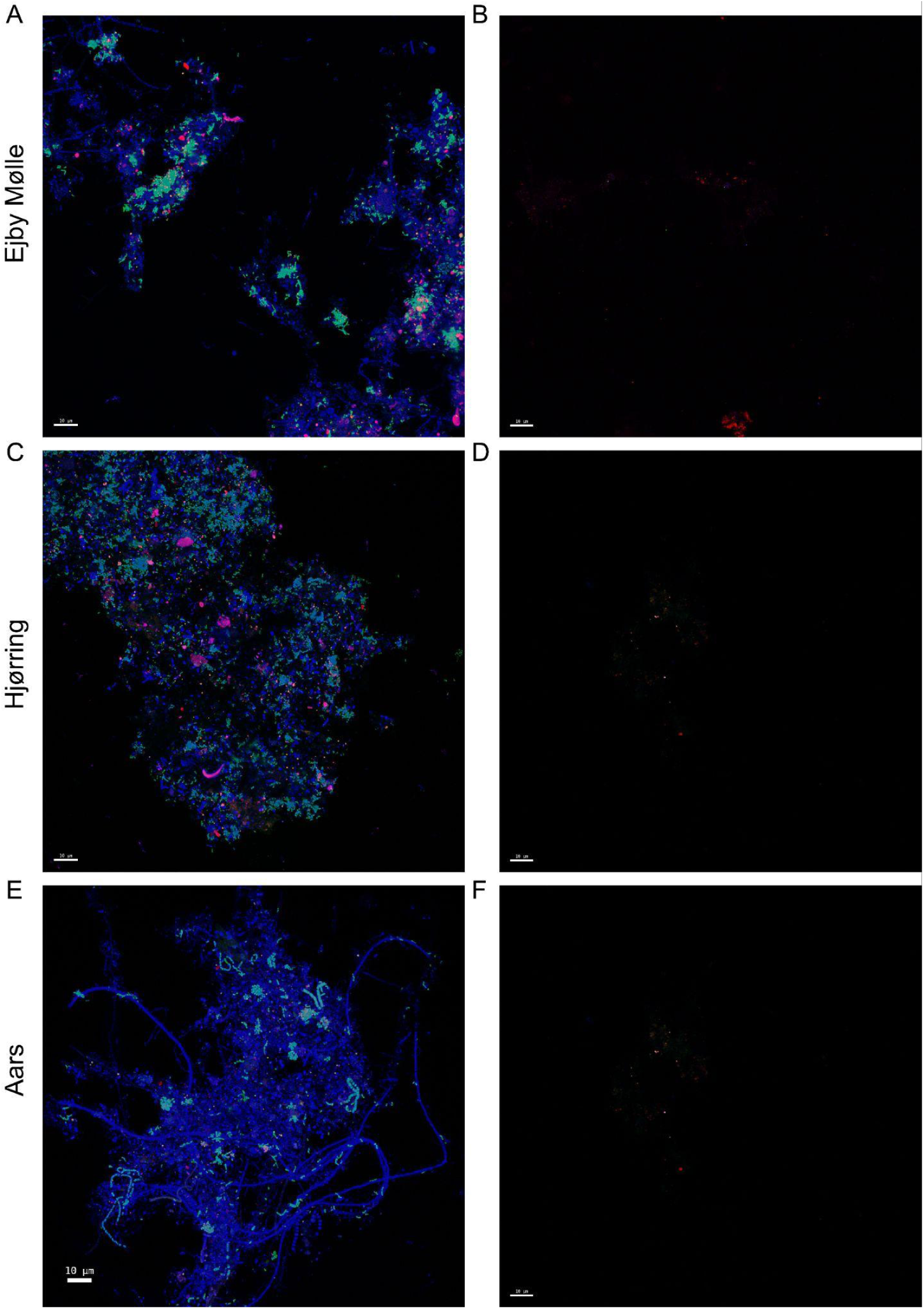
Spatial coaggregation of *Ca*. Lutisaccharimonas and *Ca.* Phosphoribacter and negative controls. Representative FISH images of activated sludge samples from the Ejby Mølle (A,B), Hjørring (C,D) and Aars (E,F) WWTPs. In panels A, C and E, *Ca.* Phosphoribacter was labeled with probe Phos601 (Atto488; green), *Ca*. Lutisaccharimonas with probe Sac1343 (Cy3; magenta) and total bacteria with the EUB338 probe mix (Cy5; blue). Negative controls using nonEUB in Atto488 (green), Cy3 (red) and Cy5 (blue) are shown in panels B, D and F. Scale bars in all panels are 10 µm.

Samples from Horsens WWTP were chosen for a quantitative spatial coaggregation analysis because of the high abundance of *Ca*. Phosphoribacter and *Ca.* Lutisaccharimonas. The spatial pair cross-correlation function of the two populations, estimated by the linear dipole method, was significantly elevated above the random expectation (y = 1, see Methods) within the short-distance range, demonstrating a positive spatial association between both populations at distances of up to approximately 20 µm (Fig. 1G,H). This coaggregation signal was strongest within less than 5 µm and progressively declined with increasing distance. Beyond approximately 20 µm, the pair cross-correlation function approached 1, indicating a random distribution of the two populations relative to each other at larger spatial scales. These results suggest a close spatial association between *Ca*. Lutisaccharimonas and *Ca.* Phosphoribacter within their direct local neighborhood, which is consistent with an epibiotic relationship.

The directionality of the spatial association between *Ca*. Lutisaccharimonas and *Ca.* Phosphoribacter was subsequently evaluated using the inflate algorithm (see Methods). For this purpose, either population was treated once as the ‘analyzed’ and once as the ‘reference’ population. In all analyses, the observed cumulative fractions exceeded those of simulated randomly distributed populations, supporting the short-range coaggregation between *Ca*. Lutisaccharimonas and *Ca.* Phosphoribacter which was also detected by the linear dipole method (see above and Fig. 1G,H). However, this spatial association was markedly asymmetric. When *Ca*. Lutisaccharimonas was treated as the analyzed population and *Ca.* Phosphoribacter as the reference population, the normalized cumulative fraction reached values above 3 at the shortest distances (Fig. 1J). By comparison, when *Ca.* Phosphoribacter was analyzed relative to *Ca*. Lutisaccharimonas, the maximum normalized cumulative fraction did not exceed 1.8 (Fig. 1I). Thus, a substantially larger proportion of *Ca*. Lutisaccharimonas occurred close to *Ca.* Phosphoribacter than expected for a random spatial distribution, whereas the reciprocal spatial enrichment of *Ca.* Phosphoribacter around *Ca*. Lutisaccharimonas was significantly weaker. This asymmetry is consistent with a *Ca.* Phosphoribacter-dependent lifestyle of *Ca*. Lutisaccharimonas (Fig. 1I–L).

A recent study found *Zoogloea* cells in close association with Patescibacteria to display a lower fluorescence intensity after FISH with rRNA-targeted probes than other *Zoogloea* and interpreted this result as an indication of reduced metabolic activity of *Zoogloea* after attachment of the Patescibacteria^19^. To investigate whether the interaction between *Ca.* Phosphoribacter and *Ca*. Lutisaccharimonas might have a similar effect detectable by FISH, we measured the fluorescence intensity of *Ca.* Phosphoribacter but found that it did not differ consistently between *Ca.* Phosphoribacter located close to and farther from *Ca*. Lutisaccharimonas. Across distance thresholds *r* of 0.5 to 3 µm, the median image-level fluorescence differences between *Ca.* Phosphoribacter at distances ≤ *r* and > *r* from *Ca*. Lutisaccharimonas ranged only from -1 to +1 intensity units, with bootstrap 95% confidence intervals spanning or reaching zero (Supplementary Fig. 3). Paired Wilcoxon signed-rank tests likewise detected no significant differences at any distance threshold (Holm-adjusted *P* values were: 1.0 for *r* = 0.5 µm; 0.99 for *r* = 1 µm; 1.0 for *r* = 2 µm; 0.64 for *r* = 3 µm).

### Co-occurrence of *Ca*. Lutisaccharimonas and *Ca.* Phosphoribacter in global and Danish WWTPs

We next asked whether the physical association between *Ca*. Lutisaccharimonas and *Ca.* Phosphoribacter observed by FISH was reflected by the population abundance distributions of the two taxa. Our previous analysis of 565 globally distributed WWTP samples had identified positive associations between *Ca.* Phosphoribacter and *Ca*. Lutisaccharimonas at both genus and ASV levels^6^ (Fig. 2A,B). To assess whether these associations persisted through time, we reanalyzed weekly or biweekly samples collected over approximately one year from three Danish EBPR plants using a modified 16S rRNA gene primer set with improved coverage of Patescibacteria^6^.

**Fig. 2.**
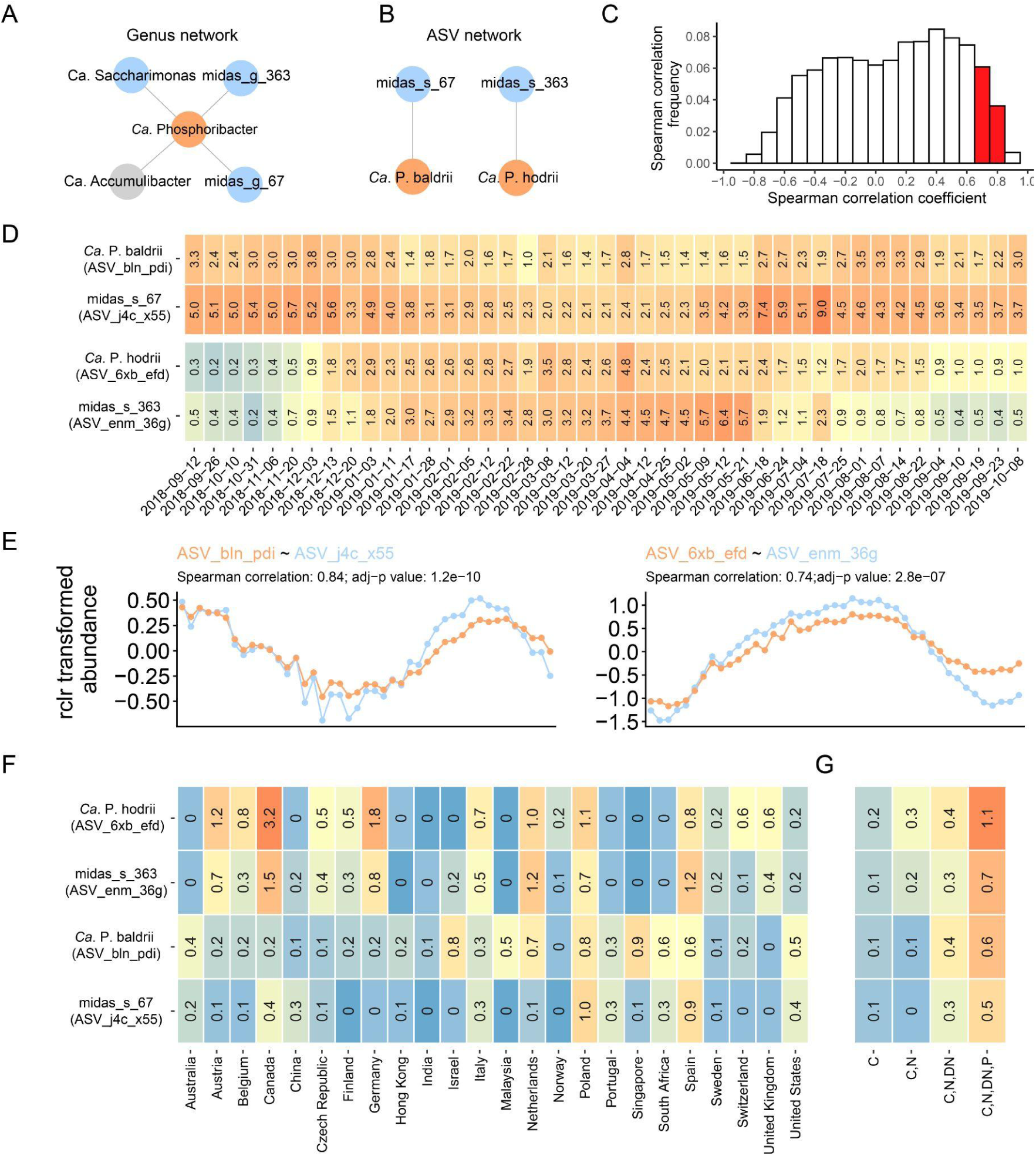
Co-occurrence analyses of *Ca*. Lutisaccharimonas and *Ca.* Phosphoribacter in the Esbjerg West WWTP. (A) Excerpt from a genus-level co-occurrence network constructed from 16S rRNA gene amplicon data from global WWTPs^6^. (B) Excerpt from an ASV-level co-occurrence network constructed from global WWTPs^6^. (C) Spearman correlation coefficient distribution histogram of 6,555 ASV pairs from the Esbjerg West WWTP dataset determined in this study. The bars representing the *Ca.* Phosphoribacter–*Ca*. Lutisaccharimonas ASV pairs are highlighted in red. (D) Relative abundance of ASVs of *Ca.* P. baldrii*, Ca.* P. hodrii, *Ca*. Lutisaccharimonas ASV_j4c_x55 and *Ca*. Lutisaccharimonas ASV_enm_36g in the Esbjerg West WWTP during the sampling period. (E) Correlation plot of the robust centered log-ratio (rclr)-transformed (to handle sparsity and compositionality) abundances of the ASV pairs of *Ca.* P. baldrii (ASV_bln_pdi) and *Ca*. Lutisaccharimonas ASV_j4c_x55 (left) and *Ca.* P. hodrii (ASV_6xb_efd) and *Ca*. Lutisaccharimonas ASV_enm_36g (right) across the sampling period in the Esbjerg West WWTP. The x-axis labeling (sampling date) is shown in panel D. (F,G) The average relative abundance of the two most abundant ASVs of *Ca*. Lutisaccharimonas and two most abundant ASVs of *Ca.* Phosphoribacter in the activated sludge samples per country (F) and per process type (G) collected from global WWTPs^6^.

At the Esbjerg West WWTP, a *Ca*. P. baldrii ASV (ASV_bln_pdi) and a *Ca*. Lutisaccharimonas ASV (ASV_j4c_x55) consistently displayed high relative abundances, together comprising up to ∼10% of the microbial community. A second *Ca*. Lutisaccharimonas ASV (ASV_enm_36g) displayed a more transient abundance pattern, with relative abundances below 1% before December 2018 and after July 2019. During the intervening period, its relative abundance was higher, reaching a maximum of 6.4% (Fig. 2D). ASV-level correlation analysis in Esbjerg West samples identified two highly correlated pairs: *Ca.* P. baldrii with *Ca*. Lutisaccharimonas ASV_j4c_x55 (r = 0.84, adjusted *P* value = 1.2×10^−10^) and *Ca.* P. hodrii with *Ca*. Lutisaccharimonas ASV_enm_36g (r = 0.74, adjusted *P* value = 2.8×10^−07^) (Fig. 2E). These correlations ranked 53^rd^ and 318^th^, respectively, among 6,555 ASV pairs (Fig. 2C). Stronger correlations were observed almost exclusively between different ASVs from the same genus, likely reflecting intragenomic heterogeneity among 16S rRNA gene copies or closely related microorganisms with similar physiological traits or environmental niches. Positive correlations between *Ca.* Phosphoribacter species and *Ca*. Lutisaccharimonas ASVs were also observed in the other two Danish WWTPs (Supplementary Figs. 4 and 5 and Supplementary notes). Thus, the same dominant *Ca*. Lutisaccharimonas and *Ca.* Phosphoribacter populations were prevalent in EBPR plants globally and exhibited strongly correlated abundance patterns across three independent Danish time series. Although correlation alone cannot establish a direct interaction, its consistency across species pairs, treatment plants and sampling scales supports a persistent ecological association and complements the physical association detected by FISH.

The *Ca*. Lutisaccharimonas ASVs ASV_enm_36g and ASV_j4c_x55 and the *Ca.* Phosphoribacter ASVs ASV_6xb_efd and ASV_bln_pdi were also the dominant representatives of their respective genera in the global activated sludge dataset and were particularly prevalent and abundant in EBPR plants. Together, these results indicate that the association detected in the Danish plants involves globally distributed populations characteristic of EBPR systems (Fig. 2F,G).

### CRISPR spacer matches and HGT events between *Ca*. Lutisaccharimonas and *Ca.* Phosphoribacter

Because close cellular associations can leave genomic signatures of DNA exchange^20,21^, we searched for independent evidence linking *Ca*. Lutisaccharimonas to *Ca.* Phosphoribacter. We first screened 22,277 high-quality bacterial and archaeal metagenome-assembled genomes (MAGs) recovered from 83 WWTPs for CRISPR spacers matching genomic sequences from other prokaryotes. In total, 108,783 spacers were identified, of which 100,730 remained after the removal of homopolymeric sequences. These spacers were matched against the complete set of high-quality MAGs using a minimum sequence identity of 90%. After excluding self-matches, matches between MAGs with the same lowest-level taxonomic classification and matches associated with predicted viral regions, 416 unique spacers remained. Of these, 44 distinct spacer–protospacer relationships connected 49 Patescibacteria MAGs with 31 bacterial MAGs belonging to seven non-Patescibacteria phyla (Fig. 3; Supplementary Table 2).

**Fig. 3.**
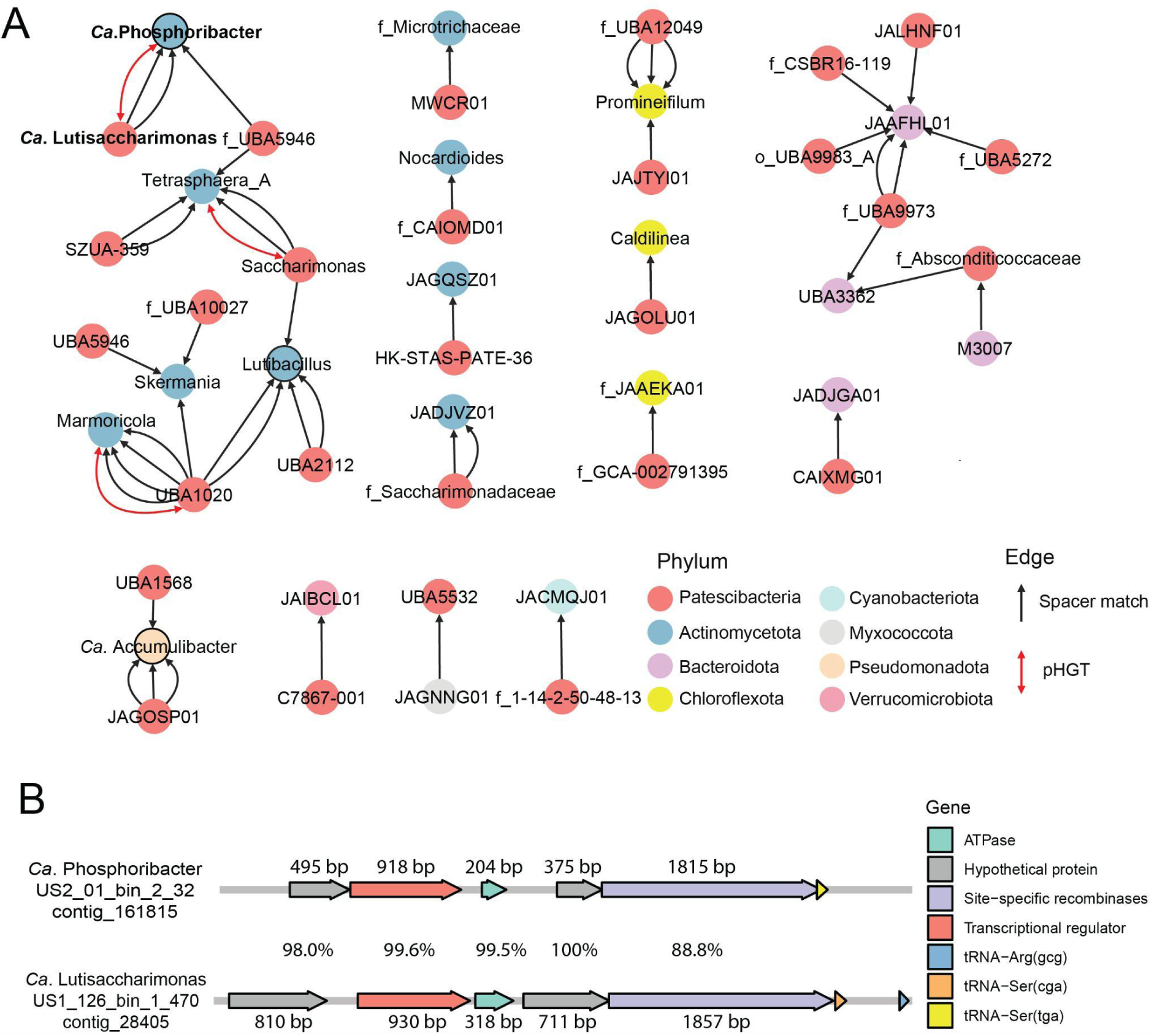
(A) CRISPR-based inferences of potential interaction partners of Patescibacteria and other WWTP microbiome members. Each node represents a genus or an unclassified genus within a family, which is either a CRISPR spacer donor or acceptor. The arrow direction indicates the potential spacer acquisition from the donor to the recipient. The number of arrows between a pair of nodes indicates the number of different spacers detected in the putative recipient genus. The genera *Ca*. Lutisaccharimonas and *Ca.* Phosphoribacter are labeled in bold font. Family names are shown with the prefix “f_” when the genus is unclassified in GTDB version r220. Colored circles with a black ring represent PAOs. Raw data of the network can be found in Supplementary Table 2. Genus pairs in this network detected with a predicted horizontal gene transfer (pHGT) are labeled with a double-headed red arrow (see next section). (B) Genes that were potentially horizontally transferred between *Ca*. Lutisaccharimonas and *Ca.* Phosphoribacter detected in a MAG pair. Genes were annotated by eggnog-mapper v2^22^ and sequence identities of aligned regions are indicated.

Actinomycetota accounted for the largest number of non-Patescibacteria MAGs containing spacers that matched Patescibacteria genomes. Specifically, 13 Actinomycetota MAGs contained 21 distinct spacers matching protospacers in 23 Saccharimonadia MAGs representing 12 genera. This pattern is consistent with previous observations that characterized Saccharimonadia as frequently associated with members of the Actinomycetota^3,7,23,24^.

Within this network, we identified spacer–protospacer matches specifically connecting *Ca.* Phosphoribacter with *Ca*. Lutisaccharimonas. Two 40-bp spacers encoded by three *Ca.* Phosphoribacter MAGs matched sequences on two *Ca*. Lutisaccharimonas contigs, with only one to three mismatches (Supplementary Table 2). To assess the taxonomic distribution of these sequences more broadly, we searched for the two spacer sequences among medium-quality MAGs and unbinned contigs from the global WWTP dataset. This analysis identified 12 additional spacer sequence-containing contigs. Three belonged to medium-quality MAGs classified as *Ca*. Lutisaccharimonas, and five unbinned contigs were taxonomically assigned to Saccharimonadia. The remaining four contigs were assigned to Actinomycetota, in which the matching sequences occurred as spacers within CRISPR arrays. The distribution and genomic context of these sequences therefore support *Ca*. Lutisaccharimonas as the source of the protospacers targeted by the *Ca.* Phosphoribacter CRISPR arrays (Supplementary Table 3).

The three *Ca.* Phosphoribacter MAGs shared more than 99.4% average nucleotide identity and contained nearly identical spacer sequences, despite being independently assembled from WWTP metagenomes originating from different countries. They also encoded a conserved type III-E CRISPR–Cas system (Supplementary Fig. 6, Supplementary Table 4 and Supplementary text). The corresponding *Ca*. Lutisaccharimonas protospacers were located within two open reading frames encoding proteins of unknown function. Both open reading frames occurred in repetitive genomic regions that were present in four to six copies across the *Ca*. Lutisaccharimonas MAGs, and the protospacers were consistently associated with a 5′-TCC-3′ flanking motif. Together, these findings indicate that closely related *Ca.* Phosphoribacter populations from geographically separated WWTPs have acquired CRISPR spacers matching sequences present in *Ca*. Lutisaccharimonas genomes.

Additional spacer–protospacer relationships suggested that associations between Patescibacteria and abundant activated sludge bacteria extend beyond the *Ca*. Lutisaccharimonas–*Ca.* Phosphoribacter pair. The globally important PAO *Ca.* Accumulibacter contained spacers matching protospacers from two Patescibacteria genera, UBA1568 and JAGOSP01, whereas another PAO *Ca*. Lutibacillus contained spacers matching three Saccharimonadia genera. Spacer matches also connected members of Patescibacteria with genera belonging to Bacteroidota and the Chloroflexota class Anaerolineae (Fig. 3A and Supplementary Table 2). These predicted relationships were broadly consistent with previously reported co-occurrence patterns in activated sludge communities⁶, suggesting that CRISPR spacer matching can recover ecologically plausible associations between Patescibacteria and potential interaction partners.

Subsequently, we tested whether close contact between *Ca*. Lutisaccharimonas and *Ca.* Phosphoribacter had also resulted in horizontal gene transfer events^25^. Using a stringent threshold of more than 99% nucleotide identity across an alignment length exceeding 500 bp, we searched the global collection of 22,277 high-quality MAGs for putatively transferred sequences between Patescibacteria and members of other bacterial phyla. This analysis identified 418 genus pairs connecting Patescibacteria with non-Patescibacteria taxa, including a significant association between *Ca*. Lutisaccharimonas and *Ca.* Phosphoribacter (Fisher’s exact test, *P* = 1.4 × 10^-3^, Supplementary Table 5). The analysis also recovered gene-transfer signals for previously reported or biologically plausible associations, including *Microsaccharimonas–Leucobacter*, *Nitrospira_A–Micavibrio_A*and *Nitrospira_A–Nitrosomonas*, supporting the ability of the approach to detect close ecological relationships^24,26^.

Examination of the genomic regions underlying the *Ca*. Lutisaccharimonas–*Ca.* Phosphoribacter signal identified a putatively transferred region containing five genes (three of which were less than 500 bp and thus filtered out in our pipeline), including a transcriptional regulator and an adjacent gene of unknown function (Fig. 3B). Both genes displayed similarly high nucleotide identities between the two genomes, suggesting that they may have been transferred together. Recombinase genes located in this region further supported its potential mobility. EggNOG-mapper assigned both proteins to orthologous groups predominantly associated with Actinomycetota, consistent with transfer from a *Ca.* Phosphoribacter-related donor to *Ca*. Lutisaccharimonas.

We performed several additional analyses to evaluate whether the apparent transfer could have resulted from assembly or binning errors. The *Ca.* Phosphoribacter contig was assigned by Kaiju^27^ to *Phycicoccus*, a member of the Actinomycetota, whereas the *Ca*. Lutisaccharimonas contig was classified more broadly as bacterial. Importantly, the *Ca*. Lutisaccharimonas contig also contained a 16S rRNA gene assigned to *Ca*. Lutisaccharimonas. In addition, BLAST searches of tRNA genes adjacent to the recombinases produced best matches consistent with the taxonomic assignments of their respective MAGs. These independent observations support a genuine horizontal transfer event, most plausibly from an Actinomycetota donor related to *Ca.* Phosphoribacter into *Ca*. Lutisaccharimonas.

Beyond this focal genus pair, two additional genus pairs were independently recovered by both the CRISPR and horizontal-gene-transfer analyses: UBA1020–*Marmoricola* and *Saccharimonas–Tetrasphaera_A* (Fig. 3A). The convergence of these independent genomic approaches supports close and potentially recurrent contact between Saccharimonadia and Actinomycetota in activated sludge communities. In particular, the detection of both *Ca.* Phosphoribacter CRISPR spacers matching *Ca*. Lutisaccharimonas sequences and a transferred genomic region shared by the two taxa provides complementary evidence that *Ca*. Lutisaccharimonas and *Ca.* Phosphoribacter have exchanged genetic material. Together with their spatial coaggregation and coordinated population dynamics, these genomic signatures strengthen the inference of a persistent association between the two lineages in geographically separated WWTPs.

### *Ca*. Lutisaccharimonas possesses an expanded metabolic repertoire

To investigate the metabolic basis of the association between *Ca*. Lutisaccharimonas and *Ca.* Phosphoribacter, we compared the genomic repertoire of *Ca*. Lutisaccharimonas with those of other Saccharimonadia. We compiled 865 high-quality Saccharimonadia MAGs from the global activated sludge dataset and 871 additional genomes from GTDB release 220^17^. A subset of 1,337 species-dereplicated MAGs was selected from this collection of 1,736 Saccharimonadia MAGs. The 77 species-dereplicated MAGs classified within the family UBA4665 were subsequently selected for pangenome analysis. Functional enrichment analysis identified 114 gene families that were significantly enriched in *Ca*. Lutisaccharimonas relative to other members of the family. These gene families were predominantly associated with nucleotide biosynthesis, amino acid metabolism and the synthesis or utilization of intracellular storage compounds (Fig. 4, Supplementary Table 6).

**Fig. 4.**
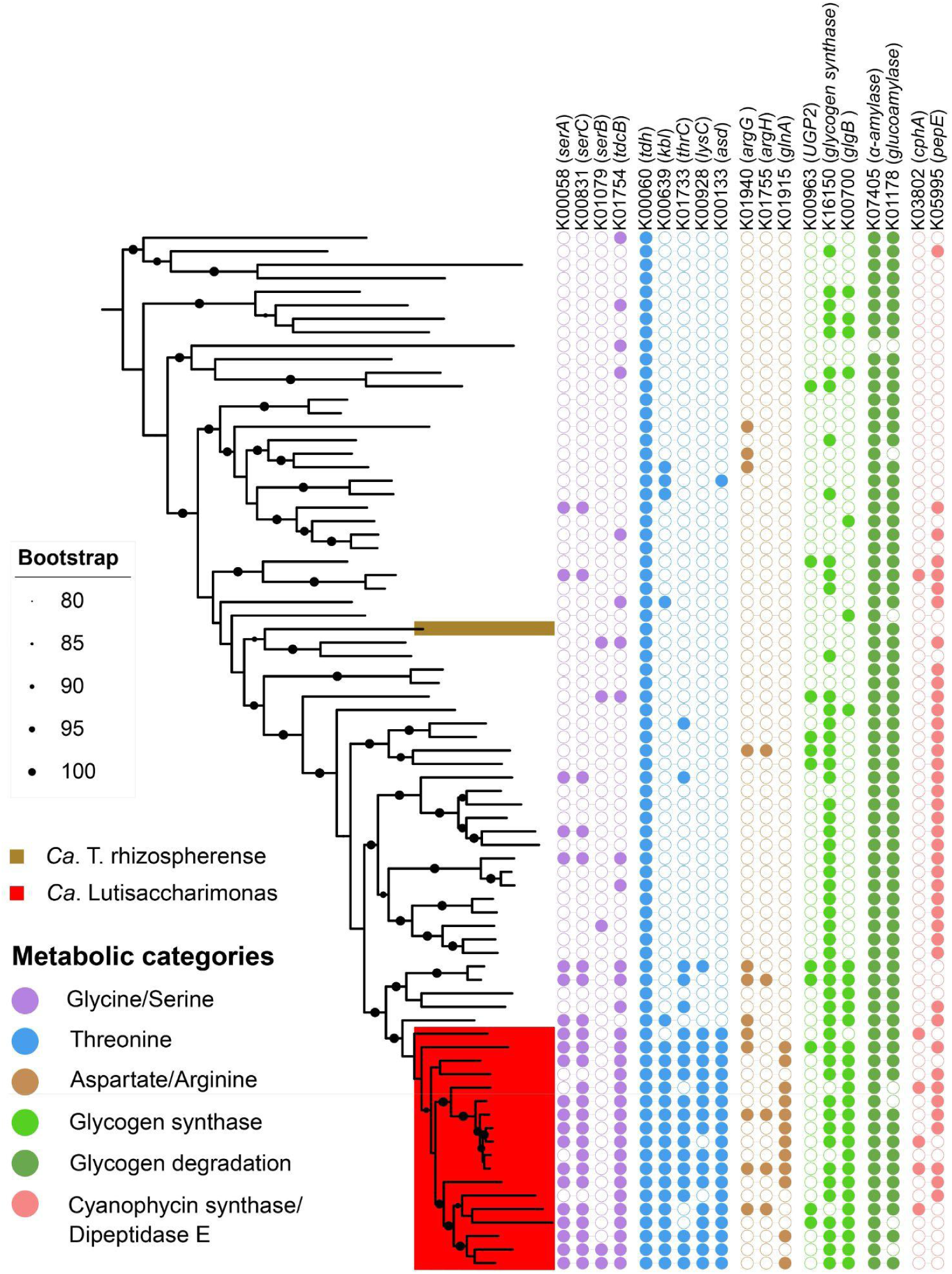
Amino acid, glycogen, and cyanophycin metabolism genes among family UBA4665 members within the Saccharimonadia. The phylogenetic tree was constructed based on GTDB r220 single-copy marker genes with IQ-TREE 2. The presence of metabolic genes is indicated with a filled circle and their absence is indicated with an empty circle. *Ca*. Lutisaccharimonas MAGs within the UBA4665 family are highlighted in red, whereas *Ca*. Teamsevenus rhizospherense29 is labeled in brown.

Most notably, all analyzed *Ca*. Lutisaccharimonas genomes encoded near-complete pathways for *de novo* nucleotide biosynthesis (Extended Data Fig. 2, Supplementary Table 7, Supplementary text). These included the KEGG modules for phosphoribosyl pyrophosphate biosynthesis (M00005), inosine monophosphate biosynthesis (M00048), adenine ribonucleotide biosynthesis (M00049), guanine ribonucleotide biosynthesis (M00050) and pyrimidine ribonucleotide biosynthesis (M00051). This capacity is uncommon among Patescibacteria, most of which lack substantial portions of the pathways required for *de novo* purine and pyrimidine biosynthesis. The conservation of these pathways across *Ca*. Lutisaccharimonas therefore indicates a greater capacity for autonomous nucleotide production than is typical of this phylum.

**Extended Data Fig. 2.**
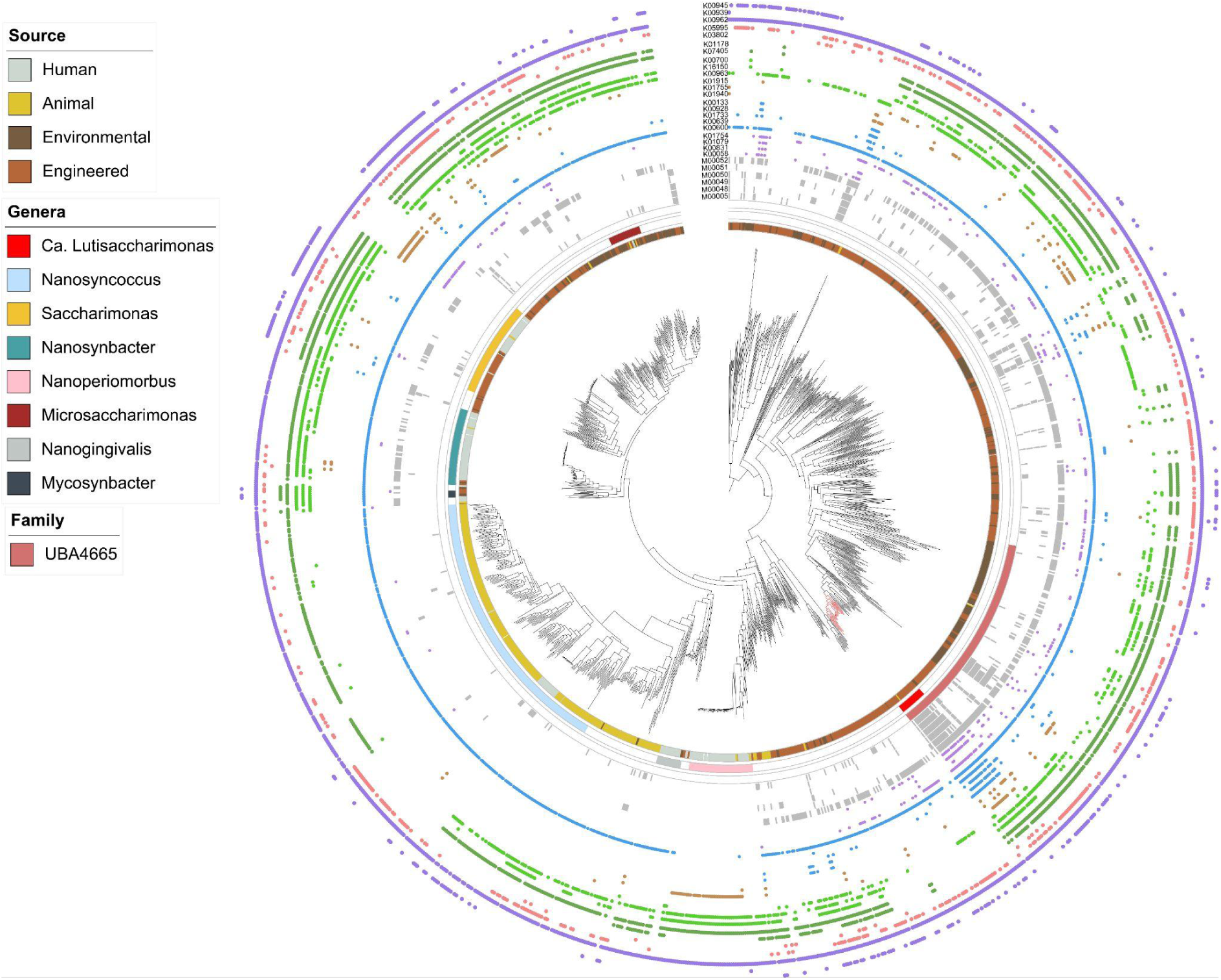
Maximum-likelihood phylogenetic tree of 1,337 species-representative MAGs of Saccharimonadia. The phylogenetic tree was constructed from the concatenated alignment of 120 single-copy marker gene proteins from GTDB-TK. The tree was inferred by IQ-TREE 2 which automatically selected the LG+F+I+R10 model. The color stripes (from inside to outside) represent (1) the type of sample source from which the organism or MAG was retrieved; (2) the genus *Ca*. Lutisaccharimonas and other genera with a proposed candidate name; (3) the family UBA4665; (4–9) the presence of genes of KEGG modules labeled in the circles with ≥75% completeness.

Despite this expanded biosynthetic potential, *Ca*. Lutisaccharimonas may have retained the ability to obtain nucleotides from external RNA. All *Ca*. Lutisaccharimonas genomes encoded polynucleotide phosphorylase (PNPase; K00962), which catalyzes phosphorolytic RNA degradation and releases nucleoside diphosphates (Extended Data Fig. 2). Furthermore, *Ca*. Lutisaccharimonas and *Ca.* Phosphoribacter displayed complementary preferences for several synonymous codons (Supplementary Fig. 7), albeit much less pronounced than for other host-Patescibacteria pairs^28^. Complementary codon usage has been proposed to limit the translation of host-derived transcripts after their acquisition by Patescibacteria, thereby facilitating the use of RNA primarily as a nucleotide source^28^. This pattern might be consistent with the lifestyle of *Ca*. Lutisaccharimonas being less dependent on an external nucleotide supply from host cells than that of most other Patescibacteria.

*Ca*. Lutisaccharimonas genomes were also enriched in genes involved in serine and threonine metabolism. These included *serA*, encoding D-3-phosphoglycerate dehydrogenase (K00058), and *serC*, encoding phosphoserine aminotransferase (K00831), which catalyzes the conversion of 3-phosphoglycerate to phosphoserine. By contrast, serB, encoding the phosphoserine phosphatase required for the final step of canonical serine biosynthesis, was detected in only two *Ca*. Lutisaccharimonas MAGs. The widespread presence of *serA* and *serC* therefore does not indicate complete serine biosynthesis but suggests that *Ca*. Lutisaccharimonas can transform intermediates of serine metabolism. *Ca*. Lutisaccharimonas genomes were additionally enriched in *tdcB*, encoding threonine dehydratase (K01754), which can convert serine or threonine to pyruvate or 2-oxobutanoate, respectively. Genes encoding threonine dehydrogenase (*tdh*; K00060) and 2-amino-3-ketobutyrate CoA ligase (*kbl*; K00639) were also enriched, providing a potential route from threonine to glycine and acetyl-CoA. Together, these genes suggest that *Ca*. Lutisaccharimonas can channel serine- and threonine-derived metabolites into its central carbon metabolism (Fig. 4).

Several genes involved in aspartate and arginine metabolism were likewise enriched in *Ca*. Lutisaccharimonas. These included *lysC*, encoding aspartate kinase (K00928), and *asd*, encoding aspartate-semialdehyde dehydrogenase (K00133), which direct aspartate-derived carbon into pathways leading to several amino acids and metabolic intermediates. The enrichment of *argG* and *argH*, encoding argininosuccinate synthase (K01940) and argininosuccinate lyase (K01755), respectively, indicated the potential to convert aspartate-derived intermediates to arginine and fumarate. Glutamine synthetase (*glnA*; K01915), which catalyzes the ATP-dependent conversion of glutamate and ammonia to glutamine, was also enriched (Fig. 4). These findings indicate that *Ca*. Lutisaccharimonas has an expanded capacity to transform amino acids and their intermediates compared with other members of the UBA4665 family.

Additional enriched functions were associated with carbohydrate and storage-polymer metabolism. A subset of the *Ca*. Lutisaccharimonas MAGs encoded UTP–glucose-1-phosphate uridylyltransferase (UGP2; K00963), glycogen synthase (K16150) and the glycogen-branching enzyme *glgB* (K00700), indicating the genomic potential for glycogen synthesis in some members of the genus. Conversely, nearly all *Ca*. Lutisaccharimonas genomes encoded glycoside hydrolases potentially involved in α-glucan degradation, including α-amylase (K07405) and glucoamylase (K01178) (Fig. 4). The broader distribution of these degradative enzymes suggests that the utilization of glycogen or related α-glucans is more widely conserved than glycogen synthesis within *Ca*. Lutisaccharimonas. Other enriched functions associated with carbohydrate, secondary-metabolite and oxygen metabolism are described in the Supplementary text and Supplementary Fig. 8.

Several of these metabolic features converge on compounds that may be synthesized or stored by *Ca.* Phosphoribacter. In particular, *Ca*. Phosphoribacter has been proposed to accumulate cyanophycin, a nitrogen- and carbon-storage polymer consisting of a polyaspartate backbone with arginine side chains^13^. Five of the 21 *Ca*. Lutisaccharimonas MAGs encoded a cyanophycin synthetase, indicating that some *Ca*. Lutisaccharimonas populations may also synthesize this polymer (Fig. 4). None of the analyzed *Ca*. Lutisaccharimonas genomes encoded a canonical cyanophycinase. However, 13 genomes contained a family S51 dipeptidase E, which could potentially hydrolyze β-aspartyl-arginine dipeptides produced during cyanophycin degradation. This distribution raises the possibility that *Ca*. Lutisaccharimonas does not directly degrade intact host cyanophycin but instead uses cyanophycin-derived dipeptides released by *Ca.* Phosphoribacter or other community members. The expanded genomic repertoire for amino acid utilization could enable *Ca*. Lutisaccharimonas to metabolize aspartate and arginine released during dipeptide hydrolysis.

Collectively, these genomic features indicate that *Ca*. Lutisaccharimonas possesses a broader metabolic repertoire than typically found in Patescibacteria. Near-complete nucleotide biosynthesis pathways may reduce its dependence on a continuous acquisition of host-derived nucleotides, potentially allowing *Ca*. Lutisaccharimonas to remain metabolically active during periods when it is not attached to a host. At the same time, incomplete amino acid biosynthetic pathways and the predicted capacity to use amino acids, α-glucans and cyanophycin-derived products suggest that *Ca*. Lutisaccharimonas remains dependent on externally supplied metabolites. Because several of these compounds are associated with the storage metabolism of *Ca*. Phosphoribacter^13^, the comparative genomic data provide a plausible nutritional framework for the observed *Ca*. Lutisaccharimonas–*Ca.* Phosphoribacter association, although direct transfer of these metabolites remains to be demonstrated experimentally.

## Discussion

Our previous co-occurrence analysis of microbial taxa in a global collection of samples from WWTPs provided the hypothesis that uncultured members of a new patescibacterial genus, which we named here *Ca*. Lutisaccharimonas, might use the important but still uncultured PAO *Ca.* Phosphoribacter as a potential host^6^. Here, we experimentally tested this hypothesis using direct microscopic, temporal, and genomic analyses to collectively demonstrate a persistent and geographically widespread epibiotic association between *Ca.* Lutisaccharimonas and *Ca.* Phosphoribacter across multiple full-scale EBPR systems.

A recent quantitative stable-isotope-probing study in full-scale EBPR samples provided further support for metabolic coupling between these organisms^30^. In the analyzed EBPR microbial community, *Ca*. Lutisaccharimonas and several other Saccharimonadia incorporated substantial amounts of carbon derived from ^13^C-labeled acetate despite lacking the genomic capacity for direct acetate assimilation. Because *Ca.* Phosphoribacter actively assimilates acetate, this observation is consistent with indirect transfer of acetate-derived carbon from *Ca.* Phosphoribacter or other primary consumers to *Ca*. Lutisaccharimonas. However, fully resolving whether this interaction is parasitic, commensal, or mutualistic will depend on future co-culture experiments and direct functional assays.

The storage physiology of *Ca*. Phosphoribacter may help explain its suitability as an interaction partner for *Ca*. Lutisaccharimonas. *Ca.* Phosphoribacter is a globally important PAO that can account for a substantial fraction of the microbial community in Danish EBPR plants^13^. Unlike the well-characterized PAO *Ca.* Accumulibacter, *Ca*. Phosphoribacter shows no detectable *in situ* accumulation of polyhydroxyalkanoates or glycogen despite encoding genes associated with glycogen synthesis^13^. Instead, it has been proposed to store carbon and energy through the accumulation of amino acids, particularly aspartate, glutamate, arginine and glycine^31–33^, as well as the amino acid polymer cyanophycin^13^. We here demonstrated that *Ca*. Lutisaccharimonas encodes pathways that could use serine, threonine, aspartate and related intermediates in central metabolism. Its capacity to degrade α-glucans and potentially hydrolyse cyanophycin-derived β-aspartyl-arginine dipeptides further suggests that compounds associated with *Ca*. Phosphoribacter storage metabolism could serve as its nutrient sources.

*Ca*. Lutisaccharimonas differs metabolically from most characterized Patescibacteria^2^. Its near-complete pathways for *de novo* purine and pyrimidine biosynthesis suggest reduced dependence on externally supplied nucleotides. At the same time, the conservation of polynucleotide phosphorylase and complementary synonymous-codon preferences between *Ca*. Lutisaccharimonas and *Ca.* Phosphoribacter suggests that *Ca*. Lutisaccharimonas may retain the capacity to supplement nucleotide biosynthesis through the degradation of externally acquired RNA. Maintaining *de novo* nucleotide synthesis could buffer *Ca*. Lutisaccharimonas against temporal variation in host-derived metabolites during the alternating anaerobic and aerobic phases of EBPR operation, potentially supporting limited metabolic activity while unattached. However, *Ca*. Lutisaccharimonas still lacks complete pathways for several amino acids and essential cellular components, including heme, quinones and specific cofactors that can be synthesized by *Ca.* Phosphoribacter. *Ca*. Lutisaccharimonas may therefore represent an intermediate form of metabolic dependence: more biosynthetically versatile than many Patescibacteria but still reliant on an associated bacterium for key metabolites.

Beyond the focal *Ca*. Lutisaccharimonas–*Ca*. Phosphoribacter pair, our CRISPR and horizontal-gene-transfer networks suggested additional associations between Patescibacteria and PAOs, including *Ca*. Accumulibacter, *Ca*. Lutibacillus and *Tetrasphaera*. Although these associations require direct experimental validation, they raise the possibility that PAOs are particularly suitable partners for metabolically streamlined bacteria in EBPR systems. PAOs repeatedly synthesize and mobilize substantial intracellular reserves, including polyphosphate, amino acids, glycogen and polyhydroxyalkanoates, in response to alternating redox conditions. These dynamic pools may provide accessible and periodically replenished nutrient sources for attached Patescibacteria. Host selection may therefore be influenced not only by phylogenetic compatibility and cell-surface properties but also by the storage physiology of potential hosts.

Collectively, our findings strongly connect a member of the Patescibacteria with a biotechnologically important PAO and provide a framework for investigating how epibionts influence EBPR communities. The consistent association between *Ca*. Lutisaccharimonas and *Ca*. Phosphoribacter suggests that interactions involving Patescibacteria may represent an overlooked component of PAO ecology.

## Methods

### FISH and image analysis

Samples for FISH analysis were collected as part of another study^18^ from Danish EBPR WWTPs located at Ejby Mølle (28 October 2008), Hjørring (29 August 2018), Horsens (19 August 2006) and Aars (19 August 2006) and fixed with 48% ethanol (final concentration). FISH was performed as previously described^34^ with the dehydration step omitted. The applied 16S rRNA-targeted oligonucleotide probes and their target organisms are listed in Supplementary Table 1. Labeled FISH probes, helper probes, and competitor oligonucleotides were added in equimolar concentrations. CloneFISH^35^ was applied to determine the optimal formamide concentrations for both newly designed probes Sac732 and Sac1343 targeting *Ca*. Lutisaccharimonas (Supplementary Fig. 9). Per-cell fluorescence intensities at different formamide concentrations were measured by digital image analysis using the software *daime*^36^ version 3.2, which was also used to determine the probe dissociation profiles from the intensity data. Clustered cells were excluded from analysis. After further validation of the probes in FISH experiments with activated sludge samples, 30% formamide concentration in the hybridization buffer was selected as the proper stringency level for probe specificity.

The EUB338 probe mix (probes EUB338 I-III) was used for total bacterial biomass detection, and probe nonEUB338 was used in negative control FISH experiments. All probes were purchased from Biomers. Hybridizations were performed overnight with 30% formamide concentration in the hybridization buffers. A Leica SP8 confocal microscope equipped with a white-light laser was used for image acquisition.

FISH images recorded for visualization purposes were deconvoluted with Huygens Essential software v26.04 using the Decon Express function with standard settings (Scientific Volume Imaging, The Netherlands). For coaggregation analyses, samples from WWTP Horsens were chosen because of the high abundance of both *Ca*. Lutisaccharimonas and *Ca.* Phosphoribacter in this system. In total 33 randomly chosen fields of view were recorded with the same microscope settings for downstream processing and analysis with the *daime* software version 3.2. Before analysis, images of *Ca.* Lutisaccharimonas were treated by local average subtraction with a kernel size of 117^2^ pixels for background reduction. The *Ca.* Phosphoribacter and *Ca.* Lutisaccharimonas images were segmented by global intensity thresholding with the Isodata algorithm, excluding small objects up to 111 or 30 pixels, respectively, which represented noise (different object size thresholds for noise were applied to account for the different sizes of the bacterial target cells). In the segmented *Ca.* Lutisaccharimonas images, the objects remaining after the first noise filtering step were sequentially processed by the following steps to exclude autofluorescent artifacts: (1) Only objects with a mean intensity ≥60 were included; (2) these objects were partitioned by K-Means++ clustering into two size bins, of which only one bin (<1091 pixels) was included; (3) of these objects, only those with a circularity ≥0.5 were included (because *Ca.* Lutisaccharimonas cells were coccoid); (4) of these objects, only those with <250 pixels were included (because *Ca.* Lutisaccharimonas cells were small). After this procedure, only small, round-shaped, FISH-positive objects that represented *Ca.* Lutisaccharimonas remained for analysis. The mean size of 621 cells from this population was measured using the object feature quantification tool of *daime*.

The linear dipole^36,37^ and Inflate algorithms^38^ as implemented in *daime* were applied to quantify the spatial association of *Ca.* Lutisaccharimonas and *Ca.* Phosphoribacter. The linear dipole algorithm estimates the pair cross-correlation function *g*(*r*) of two microbial populations, which reflects the probability of encountering any pixels that belong to one population at defined µm distances *r* from any pixels of the other population. Since *g*(*r*) relates this probability to that expected if the populations were randomly distributed, *g*(*r*)>1, *g*(*r*) = 1 and *g*(*r*)<1 indicate coaggregation, random distribution, or segregation of the two populations, respectively.

Directionality of spatial coaggregation was tested using the Inflate algorithm. In this approach, the surfaces of all objects (cells or cell clusters) of one “reference” population are computationally expanded over increasing µm distances, and the biomass fraction of a second (the “analyzed”) population encountered within each distance is quantified. In each iteration, these biomass fractions are accumulated (summarized) over the distances tested so far. The observed cumulative fraction up to any distance *r* is normalized with that obtained for a virtually randomized “analyzed” population of equivalent abundance. Consequently, normalized values above 1 indicate that the analyzed population occurs near the reference population more frequently than expected by chance. As this approach can yield different results when the analyzed and reference populations are swapped, it can detect asymmetric coaggregation patterns where one population tends to coaggregate more frequently with the other population than vice versa.

Fluorescence intensities of *Ca*. Phosphoribacter at different distances from *Ca*. Lutisaccharimonas were measured in the same image dataset as used for the spatial analyses. Since single-cell segmentation of the densely clustered *Ca*. Phosphoribacter cells was not feasible, their cell clusters were virtually divided into grids of square-shaped “pseudo-cells” with 0.5 µm side length by using the grid tessellation tool of *daime*. Using grid cells as a replacement for single-cell segmentation is an established approach in microbial biofilm image analysis^39^. For each distance threshold *r* (with *r* = 0.5, 1.0, 2.0, or 3.0 µm), the *Ca*. Phosphoribacter grid cells were then classified as being at distances ≤ *r* or > *r* from the nearest *Ca*. Lutisaccharimonas cell. The fluorescence intensity of *Ca*. Phosphoribacter biomass was then measured within every grid cell using *daime* and the data were exported for downstream analysis in R 4.6.1^40^. To avoid pseudoreplication, fluorescence intensity was summarized at the image level by calculating the median grid cell fluorescence intensity for each distance group within each of the 33 microscopic fields. Thus, for each *r*, paired image-level medians (for the ≤ *r* and > *r* grid cell groups) were obtained. These paired image-level medians were compared using two-sided Wilcoxon signed-rank tests, chosen because measurements for both grid cell groups originated from the same images and no assumption of normality was required. *P* values from the multiple distance-threshold comparisons were adjusted using the Holm method to control the family-wise error rate. For data visualization, image-specific fluorescence intensity differences were calculated as Δ*I_r_* = *I*_≤*r*_ - *I*_>*r*_ (with *I* being the median grid cell intensity). Overall intensities and differences were summarized as medians across the 33 images, and 95% confidence intervals were estimated by non-parametric bootstrap resampling of images with 10,000 replicates.

### Sample collection, amplicon sequencing, and bioinformatics

Time-series samples were collected from three different WWTPs located at Esbjerg and Fredericia (Denmark) from 2018 to 2019. DNA extraction, amplicon sequencing with a modified V4 primer, and amplicon sequence variant (ASV) calling were performed as described previously^6^. Taxonomic annotation of ASVs was performed using DADA2 using the MiDAS 5.3 database^15^ as a reference. Samples with less than 10,000 reads were discarded before further analysis. Downstream analysis was performed in R 4.1.2^40^. Robust-centered log-ratio (rclr) transformation was conducted by the decostand function from vegan 2.6-4^41^. Analysis and visualization of the amplicon sequencing data were performed by ampvis2 2.7.17^42^, ggplot2 3.5.1^43^ and dplyr 1.1.2^44^. Spearman correlation was calculated with the rcorr function from R package Hmisc 5.0.1^45^. The *P* value of Spearman correlations was adjusted for multiple testing by the Benjamini-Hochberg method. The MiDAS 5.3 database was used for probe design and probe coverage evaluation. Full-length sequences assigned to midas_f_67 were extracted and clustered using USEARCH v11.0.667^46^ at a 99% identity threshold. Representative sequences from each cluster were aligned with MAFFT v7.526^47^ using default parameters, and a 16S rRNA gene phylogenetic tree was constructed using FastTree v2.2.0^48^.

### CRISPR analysis

A total of 22,277 HQ MAGs were obtained from the global collection of 83 WWTPs^49^. CRISPR regions of 22,277 HQ MAGs were annotated by CRISPRCasFinder^50^. Only spacers with confidence level 4 were extracted for further analysis. All nucleotide sequences from the HQ MAGs were searched against a spacer database with BLASTN-short and word_size = 10. Viral regions within the MAGs were identified using VirSorter2^51^, with a score threshold of >0.95 or a hallmark-gene count of >2, and were subsequently masked to prevent viral protospacers from being incorrectly classified as MAG-derived. Homopolymer sequences and self-hits were removed from the BLAST results. Spacer-protospacer BLAST results were filtered using an identity threshold of >90%. Network visualization was done by Cytoscape^52^. Genomes of HQ MAGs were annotated by Bakta^53^ and CRISPR-Cas genes were further annotated by HHpred^54^.

### HGT analysis

A BLAST-based method^25^ was applied to detect horizontal gene transfer events. Briefly, an all-versus-all BLASTN search using coding domain sequences of 22,277 HQ MAGs generated from 83 WWTPs was performed. BLAST results were filtered using an identity threshold of >90% and hit length >500 bp. Only inter-phylum best hits were considered as putative HGT events, which means when a gene from a MAG could be better matched to other MAGs belonging to the same phylum, it was discarded^55^. Additionally, coding genes with matches at the ends of contigs and duplicated contigs were filtered out as these are likely to be false positives due to a de Bruijn graph bubble in the assembly step^55^. Significance testing of HGT between genus pairs was performed using the fisher.test() function in R and followed by Benjamini-Hochberg FDR correction.

### Comparative genomic analysis

865 HQ MAGs of Saccharimonadia from the dataset generated from the global activated sludge collection^49^ and 871 genomes from GTDB release version 220 were collected^17^. GTDB-Tk v2.4.0^17^ was used for GTDB taxonomy classification, marker gene extraction and alignment. The alignment was used as input for IQ-TREE v2.4.0^56^ for model selection and phylogenetic tree reconstruction. The tree was visualized by iTOL 7.2^57^. Pangenome analysis was performed using anvi’o v8^58^ with an MCL inflation value of 5 using KOfam annotation with the function anvi-compute-functional-enrichment-in-pan^59^. Enrichment analysis was done using Rao’s test implemented in the anvi’o program based on gene presence/absence with adjusted P < 0.05.

## Data availability

The MAGs generated from global WWTP metagenomic datasets are deposited in ENA under BioProject PRJEB83983. The amplicon sequencing datasets are deposited in NCBI under PRJNA1433364.

## Acknowledgements

We thank Bernhard Schink for helpful feedback about the nomenclature of *Ca*. Lutisaccharimonas. The computational results of this work have been achieved using the Life Science Compute Cluster (LiSC) of the University of Vienna. We thank the Joint Microbiome Facility (JMF) laboratory technician team for help with sample preparation for amplicon sequencing. This study was supported by the Wittgenstein Award of the Austrian Science Fund (FWF) [Z383-B] awarded to M.W., and by the Austrian Science Fund (FWF) Cluster of Excellence “Microbiomes drive Planetary Health’ (10.55776; COE 7; P.P., K.K., H.D. and M.W.), the Independent Research Fund Denmark (grant 2035-00360B to M.K.D.D.), and Novo Nordisk Foundation (REThiNk, grant NNF22OC0071498 to P.H.N. and M.K.D.D.).

## Author contributions

H.H., K.K., P.P., C.W.H. and M.W. designed the study. L.L., J.M.K., M.P., P.H.N. and M.K.D.D. provided samples and data. H.H. performed the bioinformatic analyses, with contributions from L.L., A.L. and D.S. H.H. and K.K. designed the probes and performed cloneFISH. M.N. performed FISH and image processing. H.H. and H.D. performed the spatial image analyses. H.H. wrote the manuscript with input from K.K., P.P., H.D. and M.W. All authors reviewed the manuscript and provided feedback.

## References

1. Brown, C. T. et al. Unusual biology across a group comprising more than 15% of domain Bacteria. Nature 523, 208–211 (2015).

2. Castelle, C. J. et al. Biosynthetic capacity, metabolic variety and unusual biology in the CPR and DPANN radiations. Nat Rev Microbiol 16, 629–645 (2018).

3. He, X. et al. Cultivation of a human-associated TM7 phylotype reveals a reduced genome and epibiotic parasitic lifestyle. Proceedings of the National Academy of Sciences 112, 244–249 (2015).

4. Dudek, N. K. et al. Novel Microbial Diversity and Functional Potential in the Marine Mammal Oral Microbiome. Current Biology 27, 3752–3762.e6 (2017).

5. Chiriac, M.-C. et al. Ecogenomics sheds light on diverse lifestyle strategies in freshwater CPR. Microbiome 10, 84 (2022).

6. Hu, H. et al. Global abundance patterns, diversity, and ecology of Patescibacteria in wastewater treatment plants. Microbiome 12, 55 (2024).

7. Batinovic, S., Rose, J. J. A., Ratcliffe, J., Seviour, R. J. & Petrovski, S. Cocultivation of an ultrasmall environmental parasitic bacterium with lytic ability against bacteria associated with wastewater foams. Nat Microbiol 6, 703–711 (2021).

8. Zhong, Q. et al. Episymbiotic Saccharibacteria TM7x modulates the susceptibility of its host bacteria to phage infection and promotes their coexistence. Proceedings of the National Academy of Sciences 121, e2319790121 (2024).

9. Srinivas, P., Peterson, S. B., Gallagher, L. A., Wang, Y. & Mougous, J. D. Beyond genomics in Patescibacteria: A trove of unexplored biology packed into ultrasmall bacteria. Proceedings of the National Academy of Sciences 121, e2419369121 (2024).

10. Bor, B. et al. Rapid evolution of decreased host susceptibility drives a stable relationship between ultrasmall parasite TM7x and its bacterial host. Proceedings of the National Academy of Sciences 115, 12277–12282 (2018).

11. Utter, D. R., He, X., Cavanaugh, C. M., McLean, J. S. & Bor, B. The saccharibacterium TM7x elicits differential responses across its host range. ISME J 14, 3054–3067 (2020).

12. Tian, J. et al. Acquisition of the arginine deiminase system benefits epiparasitic Saccharibacteria and their host bacteria in a mammalian niche environment. Proceedings of the National Academy of Sciences 119, e2114909119 (2022).

13. Singleton, C. M. et al. The novel genus, ‘Candidatus Phosphoribacter’, previously identified as Tetrasphaera, is the dominant polyphosphate accumulating lineage in EBPR wastewater treatment plants worldwide. ISME J 16, 1605–1616 (2022).

14. Oehmen, A. et al. Advances in enhanced biological phosphorus removal: From micro to macro scale. Water Research 41, 2271–2300 (2007).

15. Dueholm, M. K. D. et al. MiDAS 4: A global catalogue of full-length 16S rRNA gene sequences and taxonomy for studies of bacterial communities in wastewater treatment plants. Nat Commun 13, 1908 (2022).

16. Zheng, H. et al. Intralineage Diversity and Global Biogeography of Ca. Phosphoribacter. Environ. Sci. Technol. 60, 15964–15976 (2026).

17. Parks, D. H. et al. GTDB: an ongoing census of bacterial and archaeal diversity through a phylogenetically consistent, rank normalized and complete genome-based taxonomy. Nucleic Acids Research 50, D785–D794 (2022).

18. Nierychlo, M. et al. MiDAS 3: an ecosystem-specific reference database, taxonomy and knowledge platform for activated sludge and anaerobic digesters reveals species-level microbiome composition of activated sludge. Water Research 182, 115955 (2020).

19. Fujii, N., et al. Unique episymbiotic relationship between Candidatus Patescibacteria and Zoogloea in activated sludge flocs at a municipal wastewater treatment plant. Environmental Microbiology Reports 16, e70007 (2024).

20. Esser, S. P. et al. A predicted CRISPR-mediated symbiosis between uncultivated archaea. Nat Microbiol 8, 1619–1633 (2023).

21. Sures, K. et al. Acquisition of Spacers from Foreign Prokaryotic Genomes by CRISPR-Cas Systems in Natural Environments. Genome Biol Evol 17, evaf201 (2025).

22. Cantalapiedra, C. P., Hernández-Plaza, A., Letunic, I., Bork, P. & Huerta-Cepas, J. eggNOG-mapper v2: Functional Annotation, Orthology Assignments, and Domain Prediction at the Metagenomic Scale. Mol Biol Evol 38, 5825–5829 (2021).

23. Wang, Y. et al. Genetic manipulation of Patescibacteria provides mechanistic insights into microbial dark matter and the epibiotic lifestyle. Cell 186, 4803–4817.e13 (2023).

24. Xie, B. et al. Type IV pili trigger episymbiotic association of Saccharibacteria with its bacterial host. Proc Natl Acad Sci U S A 119, e2215990119 (2022).

25. Smillie, C. S. et al. Ecology drives a global network of gene exchange connecting the human microbiome. Nature 480, 241–244 (2011).

26. Dolinšek, J., Lagkouvardos, I., Wanek, W., Wagner, M. & Daims, H. Interactions of nitrifying bacteria and heterotrophs: identification of a Micavibrio-like putative predator of Nitrospira spp. Applied and Environmental Microbiology 79, 2027–2037 (2013).

27. Menzel, P., Ng, K. L. & Krogh, A. Fast and sensitive taxonomic classification for metagenomics with Kaiju. Nat Commun 7, 11257 (2016).

28. Katayama, T. et al. A representative of a ubiquitous bacterial lineage parasitically feeds on host RNA. 2026.06.21.733656 Preprint at 10.64898/2026.06.21.733656 (2026).

29. Starr, E. P. et al. Stable isotope informed genome-resolved metagenomics reveals that Saccharibacteria utilize microbially-processed plant-derived carbon. Microbiome 6, 122 (2018).

30. Sampara, P., Tomatsu, A., Malmstrom, R. R. & Ziels, R. M. Quantitative DNA Stable Isotope Probing Identifies Active Microorganisms Assimilating Volatile Fatty Acids in Full-Scale Enhanced Biological Phosphorus Removal Processes. Environ. Sci. Technol. 60, 5570–5583 (2026).

31. Nguyen, H. T. T., Kristiansen, R., Vestergaard, M., Wimmer, R. & Nielsen, P. H. Intracellular accumulation of glycine in polyphosphate-accumulating organisms in activated sludge, a novel storage mechanism under dynamic anaerobic-aerobic conditions. Applied and Environmental Microbiology 81, 4809–4818 (2015).

32. Close, K. et al. The storage compounds associated with Tetrasphaera PAO metabolism and the relationship between diversity and P removal. Water Research 204, 117621 (2021).

33. Thomson, R., Close, K., Riley, A., Batstone, D. J. & Oehmen, A. Metabolic modelling of anaerobic amino acid uptake and storage by fermentative polyphosphate accumulating organisms. Water Research 280, 123512 (2025).

34. Daims, H., Stoecker, K. & Wagner, M. Fluorescence in situ hybridization for the detection of prokaryotes. in Molecular microbial ecology 208–228 (Taylor & Francis, 2004).

35. Schramm, A., Fuchs, B. M., Nielsen, J. L., Tonolla, M. & Stahl, D. A. Fluorescence in situ hybridization of 16S rRNA gene clones (Clone-FISH) for probe validation and screening of clone libraries. Environmental Microbiology 4, 713–720 (2002).

36. Daims, H., Lücker, S. & Wagner, M. Daime, a novel image analysis program for microbial ecology and biofilm research. Environmental microbiology 8, 200–213 (2006).

37. Reed & Howard. Stereological estimation of covariance using linear dipole probes. Journal of microscopy 195, 96–103 (1999).

38. Daims, H. & Wagner, M. In situ techniques and digital image analysis methods for quantifying spatial localization patterns of nitrifiers and other microorganisms in biofilm and flocs. in Methods in enzymology vol. 496 185–215 (Elsevier, 2011).

39. Hartmann, R. et al. Quantitative image analysis of microbial communities with BiofilmQ. Nature microbiology 6, 151–156 (2021).

40. R Core Team. R: A Language and Environment for Statistical Computing. (R Foundation for Statistical Computing, Vienna, Austria, 2021).

41. Oksanen, J., et al. vegan (v2. 6-4). R. (2022).

42. Andersen, K. S., Kirkegaard, R. H., Karst, S. M. & Albertsen, M. ampvis2: an R package to analyse and visualise 16S rRNA amplicon data. 299537 Preprint at 10.1101/299537 (2018).

43. Wickham, H. Ggplot2: Elegant Graphics for Data Analysis. (Springer-Verlag New York, 2016).

44. Wickham, H., François, R., Henry, L., Müller, K. & Vaughan, D. Dplyr: A Grammar of Data Manipulation. (2023).

45. Jr, F. E. H. Hmisc: Harrell Miscellaneous. (2023).

46. Edgar, R. C. Search and clustering orders of magnitude faster than BLAST. Bioinformatics 26, 2460–2461 (2010).

47. Katoh, K. & Standley, D. M. MAFFT Multiple Sequence Alignment Software Version 7: Improvements in Performance and Usability. Mol Biol Evol 30, 772–780 (2013).

48. Price, M. N., Dehal, P. S. & Arkin, A. P. FastTree 2 – Approximately Maximum-Likelihood Trees for Large Alignments. PLOS ONE 5, e9490 (2010).

49. Liu, L. et al. The MiDAS global genome catalog: 53,501 long-read MAGs representing all core prokaryotic genera in the global activated sludge microbiome. 2026.07.10.737647 Preprint at 10.64898/2026.07.10.737647 (2026).

50. Couvin, D. et al. CRISPRCasFinder, an update of CRISRFinder, includes a portable version, enhanced performance and integrates search for Cas proteins. Nucleic Acids Research 46, W246–W251 (2018).

51. Guo, J. et al. VirSorter2: a multi-classifier, expert-guided approach to detect diverse DNA and RNA viruses. Microbiome 9, 37 (2021).

52. Shannon, P. et al. Cytoscape: A Software Environment for Integrated Models of Biomolecular Interaction Networks. Genome Res 13, 2498–2504 (2003).

53. Schwengers, O. et al. Bakta: rapid and standardized annotation of bacterial genomes via alignment-free sequence identification. Microbial Genomics 7, 000685 (2021).

54. Söding, J., Biegert, A. & Lupas, A. N. The HHpred interactive server for protein homology detection and structure prediction. Nucleic Acids Res 33, W244–W248 (2005).

55. Song, W., Wemheuer, B., Zhang, S., Steensen, K. & Thomas, T. MetaCHIP: community-level horizontal gene transfer identification through the combination of best-match and phylogenetic approaches. Microbiome 7, 36 (2019).

56. Minh, B. Q. et al. IQ-TREE 2: New Models and Efficient Methods for Phylogenetic Inference in the Genomic Era. Molecular Biology and Evolution 37, 1530–1534 (2020).

57. Letunic, I. & Bork, P. Interactive Tree of Life (iTOL) v6: recent updates to the phylogenetic tree display and annotation tool. Nucleic Acids Research 52, W78–W82 (2024).

58. Eren, A. M. et al. Community-led, integrated, reproducible multi-omics with anvi’o. Nat Microbiol 6, 3–6 (2021).

59. Shaiber, A. et al. Functional and genetic markers of niche partitioning among enigmatic members of the human oral microbiome. Genome Biology 21, 292 (2020).

