## Supplementary Text and Figures for "A Saccharimonadia epibiont interacts with the polyphosphate-accumulating bacterium *Candidatus* Phosphoribacter in wastewater treatment plants"

### Supplementary Notes

#### Co-occurrence of *Ca*. Lutisaccharimonas and *Ca*. Phosphoribacter in two additional Danish WWTPs

In the Esbjerg East WWTP, *Ca.* Phosphoribacter and the *Ca*. Lutisaccharimonas ASVs were generally less abundant (0.3–2.4%) than in Esbjerg West (0.2–9%). In the Esbjerg East WWTP, the ASV pair *Ca*. P. baldrii (ASV_bln_pdi) and *Ca.* Lutisaccharimonas (ASV_j4c_x55) was more abundant and correlated positively, although barely not statistically significant (Spearman correlation = 0.50, adjusted P−value= 0.052), while no correlation was observed for the less abundant ASV pair *Ca*. P. hodrii (ASV_6xb_efd) and *Ca.* Lutisaccharimonas (ASV_enm_36g) (Spearman correlation = 0.18, adjusted P value = 0.57) (Fig. S4).

In the Fredericia WWTP, which receives significant loads of industrial sewage in addition to municipal sewage, the relative abundances of both *Ca.* Phosphoribacter and *Ca.* Lutisaccharimonas members were highly dynamic. The *Ca*. P. hodrii ASV_6xb_efd was very abundant in the first three samples from March 2018 to April 2018, accounting for up to 20% of the total microbial community. Its relative abundance decreased to 0.7–2.9% in the winter of 2018. The *Ca*. Lutisaccharimonas ASV_enm_36g showed a similar pattern as *Ca*. P. hodrii during the sampling period, while the relative abundance of *Ca*. Lutisaccharimonas ASV_j4c_x55 was low in the first three samples from November and December 2018 and then increased strongly to 9–13% in the winter of 2018. Consistent with this, and contrary to Esbjerg East, in Fredericia, the *Ca*. P. hodrii ASV_6xb_efd and the most abundant ASV of the genus *Ca*. Lutisaccharimonas ASV_enm_36g showed a significant strongly positive correlation (Spearman correlation = 0.64, adjusted P value = 0.006), but for the ASV pair *Ca*. P. baldrii ASV_bln_pdi and *Ca*. Lutisaccharimonas ASV_j4c_x55, no statistically significant correlation was detected (Fig. S5).

**_Ca._ Phosphoribacter uses a type III-E CRISPR-Cas system to acquire spacer sequences from _Ca_. Lutisaccharimonas**

To determine which type of CRISPR system was used by Ca. Phosphoribacter to acquire the spacers from *Ca*. Lutisaccharimonas, we annotated the genomic neighborhood of the CRISPR arrays of the three *Ca.* Phosphoribacter MAGs containing spacers matching putative protospacers on MAGs of *Ca*. Lutisaccharimonas. This revealed the presence of class 1, type III-E CRISPR-Cas genes, conserved among all three *Ca.* Phosphoribacter MAGs (Fig. S6). The type III-E CRISPR system was recently discovered[1]. To perform site-specific RNA cleavage, it utilizes a unique, single-protein effector complex, Cas7-11 (also known as gRAMP), which originates from the fusion of a putative Cas11 domain and multiple Cas7 subunits[1,2]. As type III-E CRISPR systems mediate RNA targeting through the Cas7-11 effector complex, the detected spacer–protospacer matches may reflect interactions with actively expressed transcripts in *Ca*. Lutisaccharimonas rather than solely DNA-level interference.

In the type III-E system of the three *Ca.* Phosphoribacter MAGs, a Cas2 protein, followed by a Cas6-RT-Cas1 protein was annotated as an adaptation module. The Cas6-RT-Cas1 domain matched a characterized protein with 100% probability (23% identity) to a characterized protein[3], which is able to acquire CRISPR spacers directly from both DNA and RNA. Downstream of Cas6-RT-Cas1 and a gene annotated as Cas6, a polo domain and a TIR-SAVED effector was annotated (98% probability; 11% identity), which has been shown to hydrolyse $NAD^+$ to cause cell death once activated by small oligonucleotide signal molecules (e.g., cyclic nucleotides)[4]. In the *Ca.* Phosphoribacter MAGs, the effector module is a gRAMP complex, consisting of a Cas7-2x protein (which two Cas7 domains are fused together), a Cas7-Cas5-Cas11 protein, a Csm3 protein, a Cas10 protein and a Cas7-Cas11 protein (Fig. S6; Supplementary Table 3). Together, these features suggest that the *Ca.* Phosphoribacter CRISPR system is capable of acquiring spacers from actively expressed genes and mounting a potent immune response involving NAD+ depletion and programmed cell death.

**CRISPR-spacer interaction between _Ca_. Phosphoribacter and other bacteria**

We also explored spacers from the CRISPR array of the *Ca.* Phosphoribacter MAGs that did not match Patescibacteria MAGs but could be mapped to non-patescibacterial MAGs from the global WWTP dataset. We identified 14 additional unique spacers in the CRISPR arrays of the *Ca.* Phosphoribacter MAGs that matched putative protospacers in 11 MAGs belonging to Actinomycetota, Bacteroidota, Myxococcota and Pseudomonadota. Strikingly, two *Ca.* Phosphoribacter (020-NO1-03.bin.1.471 and 039-SE-29.bin.1.184) MAGs recovered from WWTPs located at Norway and Sweden, respectively, shared 98% average nucleotide identity and contained identical CRISPR arrays with the same direct repeat sequences and 30 spacer sequences. Among these 30 spacers, eight were mapped with 91-100% identity to a *Myxococcota* MAG (038-SE-10.bin.1.350) (Supplementary Table 8). However, these two

*Ca.* Phosphoribacter MAGs, with CheckM2-estimated completeness values of 96.6% and 99.1%, encoded no Cas proteins, suggesting that the *cas* genes had been lost or that the CRISPR array had been transferred by mobile genetic elements[5].

**Spacers found in Patescibacteria can be mapped to other predatory bacteria**

Although CRISPR systems are rarely detected in Patescibacteria[6–8], we identified two Patescibacteria MAGs belonging to class JAEDAM01 that encoded CRISPR systems with spacers matching protospacers in the genus M3007 (family Saprospiraceae) and an unnamed genus from the phylum Myxococcota. As several cultured Saprospiraceae and Myxococcota genera are well known for bacterial predation, this suggests that at least some Patescibacteria in WWTPs are attacked by predators (Fig. 3A; Supplementary Table 2).

**Nucleotide *de novo* biosynthesis pathways**

Nucleotide *de novo* biosynthesis pathways were significantly enriched in the genus *Ca*. Lutisaccharimonas compared to other members of the family UBA4665. Notably, the gene encoding ribose-5-phosphate pyrophosphokinase (PRPS, K00948), which converts ribose-5-phosphate to 5-phospho-alpha-D-ribose-1-diphosphate (PRPP), the activated ribose donor used to build all nucleotides, was present in 88.9% of the *Ca*. Lutisaccharimonas MAGs, but was only present in 23.7% of other UBA4665 representatives (Supplementary Table 7).

The *de novo* purine biosynthesis pathway (KEGG Module M00048), responsible for converting PRPP to inosine monophosphate (IMP), was largely complete in *Ca*. Lutisaccharimonas MAGs, with the exception of genes for folate-dependent formyltransferase (*purN*, K11175 or *purT*, K08289), which were only present in 27.8% of the *Ca*. Lutisaccharimonas MAGs. In contrast, this pathway was only present in 18% of the other UBA4665 genomes and most other classes of Patescibacteria[9] (Supplementary Table 7). The folate biosynthesis pathway is mostly missing in the Patescibacteria genomes, including *Ca*. Lutisaccharimonas, but several genes involved in the bioactive folate-derivative synthesis pathway are present in the *Ca*. Lutisaccharimonas genomes and other Patescibacteria[10], including dihydrofolate reductase (*folA*, K00287), serine hydroxymethyltransferase (*glyA*, K00600), and methylenetetrahydrofolate reductase (*folD*, K01491). Together, these enzymes may support the interconversion of bioactive tetrahydrofolate derivatives required for *de novo* nucleotide biosynthesis.

Genes involved in adenine and guanine ribonucleotide biosynthesis pathways (M00049 and M00050) were also enriched in the *Ca*. Lutisaccharimonas genus (both present in all MAGs), while being less prevalent in other UBA4665 genomes (15.3% and 30.5%, respectively) (Supplementary Table 7). These include *purA* (adenylosuccinate synthetase, K01939), *purB* (adenylosuccinate lyase, K01756), *IMPDH* (IMP dehydrogenase, K00088), and *guaA* (GMP synthase, K01951). However, *adk* (adenylate kinase, K00939), which converts AMP to ADP, was conspicuously absent in all Saccharimonadia genomes, including those of *Ca*. Lutisaccharimonas. Conversely, *gmk* (guanylate kinase, K00942), responsible for converting GMP to GDP, is encoded by 44.4% of the *Ca.* Lutisaccharimonas MAGs. Genes encoding nucleoside diphosphate kinases (*ndk*, K00940), which interconvert ADP/GDP to ATP/GTP, are widely present across all Saccharimonadia genomes (Supplementary Table 7).

*De novo* pyrimidine biosynthesis pathways (M00051) (with >75% pathway completeness) were present in 88.9% of the *Ca*. Lutisaccharimonas MAGs while being only detected in 10.2% of the other UBA4665 MAGs. This pathway included genes of *carA* (K01956), *carB* (K01955), *pyrB* (K00609), *pyrC* (K01465), *pyrE* (K00762), and *pyrF* (K01591) that were present in most members of the genus *Ca*. Lutisaccharimonas, although *pyrI* (K00610, the aspartate carbamoyltransferase regulatory subunit) was consistently missing while the catalytic subunit *pyrB* (K00609) was present in 83.3% of *Ca*. Lutisaccharimonas MAGs. Genes *pyrH* (K09903), *ndk* (K00940), and *pyrG* (K01937) involved in the conversion of UMP to UDP/UTP and CDP/CTP (M00052) were more broadly distributed among Saccharimonadia genomes. Pathways for deoxyribonucleotide biosynthesis (M00053) were generally complete across all Saccharimonadia genomes (Supplementary Table 7), while the pyrimidine deoxyribonucleotide biosynthesis pathway (M00938) was generally absent from *Ca.* Lutisaccharimonas genomes, and present in only 10% of other UBA4665 family members.

#### Carbohydrate, secondary metabolite, and oxygen metabolism

Glycolysis and pentose phosphate pathways are nearly complete in members of the genus *Ca.* Lutisaccharimonas, as in other environmental Saccharimonadia[11,12]. The enrichment of trehalose-6-phosphate synthase/phosphatase genes (TPS/TPP, K16055) in *Ca*. Lutisaccharimonas suggests that trehalose may serve as an important stress protectant or carbon source. Riboflavin kinase (*ribF*, K11753) and FAD synthetase (K14656) genes were also enriched in *Ca*. Lutisaccharimonas, but were absent in other Saccharimonadia. This indicates a potential pathway for the conversion of externally derived riboflavin to flavin mononucleotide (FMN) and subsequently to flavin adenine dinucleotide (FAD) in *Ca*. Lutisaccharimonas, reducing its metabolic dependency. Furthermore, most *Ca*. Lutisaccharimonas members encode the genes for glycogen synthesis and degradation (Fig. 4).

Additionally, all subunits I–IV of cytochrome $bo_3$-type ubiquinol oxidase (*cyoABCD* with CyoB possessing all the residues to bind heme and Cu) are encoded in 88.9% of the available MAGs of *Ca*. Lutisaccharimonas (Fig. S8), flanked by genes encoding a FAD-binding oxidoreductase, a NADH dehydrogenase and a F-type ATP synthase, suggesting that members of *Ca.* Lutisaccharimonas have some capacity to utilize oxygen in membrane-associated redox processes and may conserve energy from this activity, like previously postulated for soil-dwelling members of this clade[12].However, except for *gltX* (K01885), which participates in the first step of tetrapyrrole biosynthesis, the remaining heme-biosynthesis genes and all quinone-biosynthesis genes appeared to be absent.

### Supplementary Figures

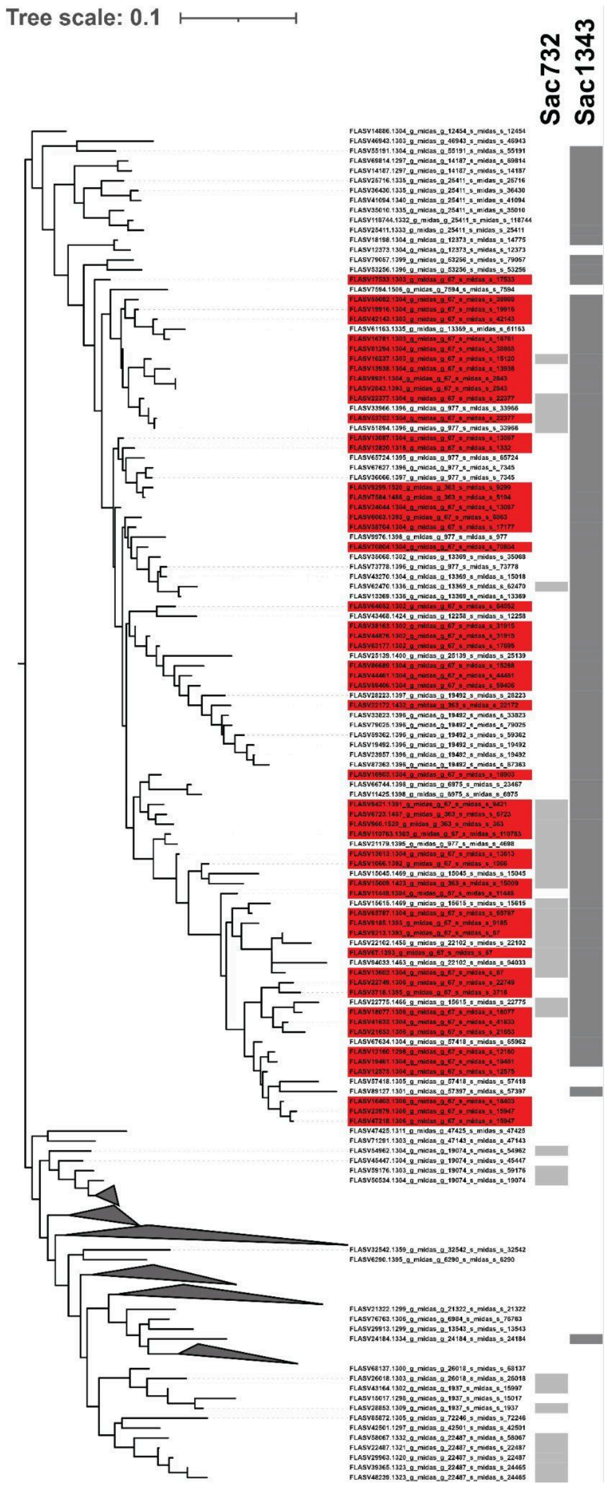


**Supplementary Fig. 1. Phylogenetic tree of full-length 16S rRNA gene sequences affiliated with midas_f_67 in the MiDAS 5.3 database and clustered at 99% sequence identity.** Sequences assigned to midas_g_67 and midas_g_363 are labeled in red, whereas the coverage of the Sac732 and Sac1343 probes is indicated by gray strips. Clades that are neither assigned to midas_g_67 nor midas_g_363 nor targeted by either probe are collapsed.

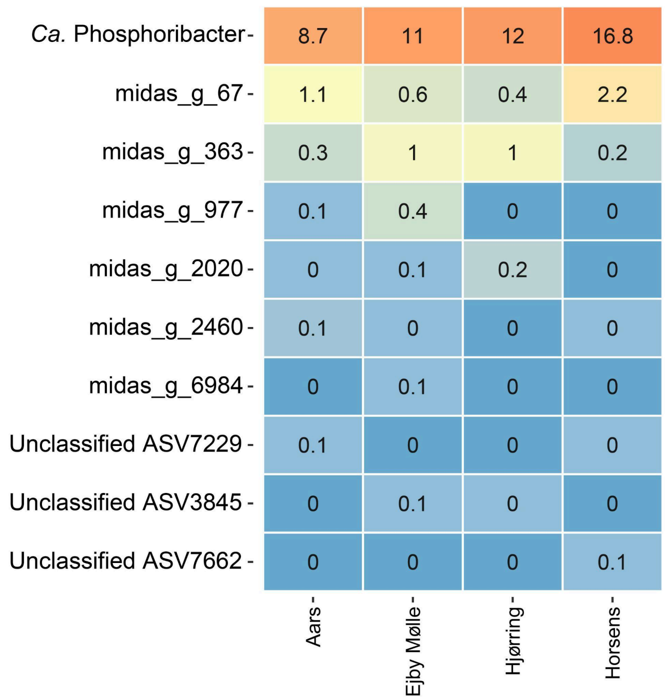


**Supplementary Fig. 2. 16S rRNA gene relative abundance data of genus *Ca.* Phosphoribacter and all genera of midas_f_67 in samples from the four analyzed Danish WWTPs.** Numbers are percent relative abundance obtained by amplicon sequencing. Data are from a previously published study[13].

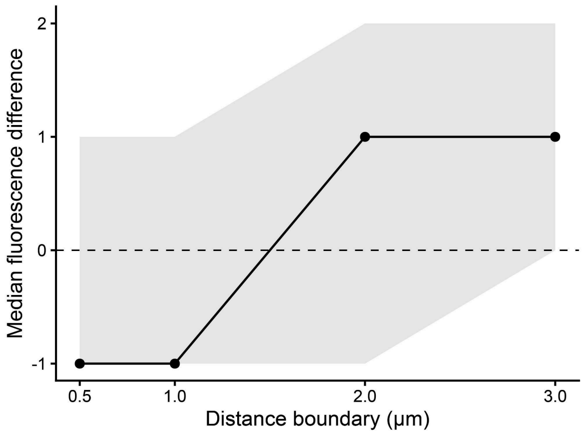


**Supplementary Fig. 3. Fluorescence intensity differences of *Ca*. Phosphoribacter across distance boundaries to *Ca*. Lutisaccharimonas.** Median differences in fluorescence intensity between *Ca*. Phosphoribacter located at distances ≤ *r* and > *r* from *Ca*. Lutisaccharimonas were calculated separately for each microscopic field and summarized across 33 fields. Points show the median paired difference ($\Delta I_r = I_{\leq r} - I_{>r}$) and the gray band indicates the bootstrap 95% confidence interval based on 10,000 image-level resamples. The dashed horizontal line denotes no difference ($\Delta I_r = 0$). Positive values indicate higher fluorescence intensity for *Ca*. Phosphoribacter closer to *Ca*. Lutisaccharimonas, and negative values indicate higher intensity for more distant *Ca*. Phosphoribacter.

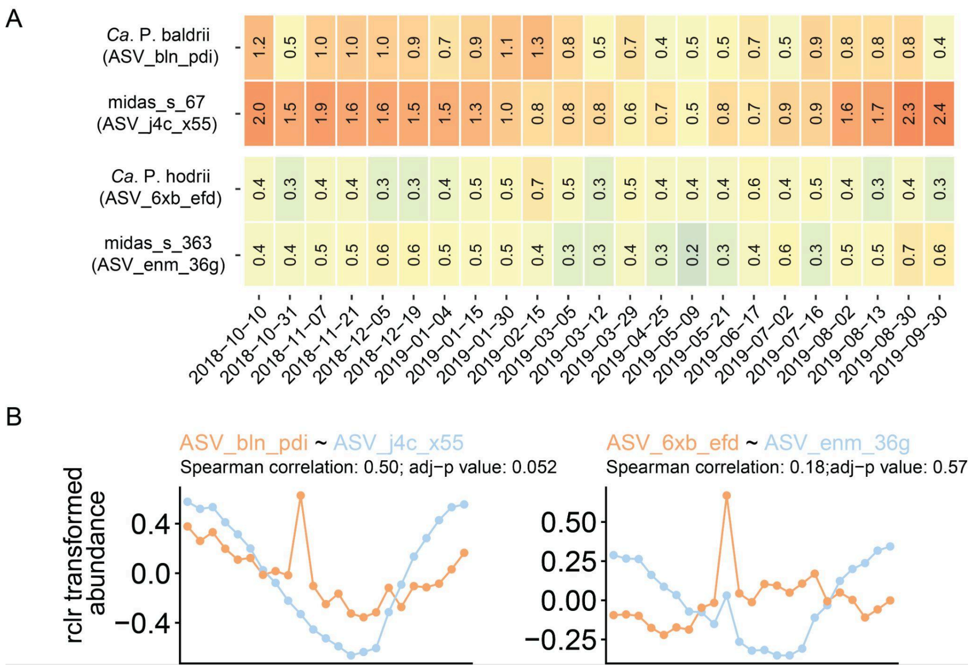


**Supplementary Fig. 4. Co-occurrence analyses of *Ca.* Lutisaccharimonas and *Ca.* Phosphoribacter in the Esbjerg East WWTP.** (A) Relative abundance of ASVs of *Ca*. P. baldrii, *Ca*. P. hodrii, *Ca*. Lutisaccharimonas ASV_j4c_x55 and *Ca*. Lutisaccharimonas ASV_enm_36g during the sampling period. (B) Correlation plot of the rclr-transformed abundance of the ASV pairs of *Ca*. P. baldrii (ASV_bln_pdi) and *Ca.* Lutisaccharimonas ASV_j4c_x55 (left), *Ca*. P. hodrii and *Ca.* Lutisaccharimonas ASV_enm_36g (right) across the sampling period. The x-axis (sampling date) is shown in panel A.

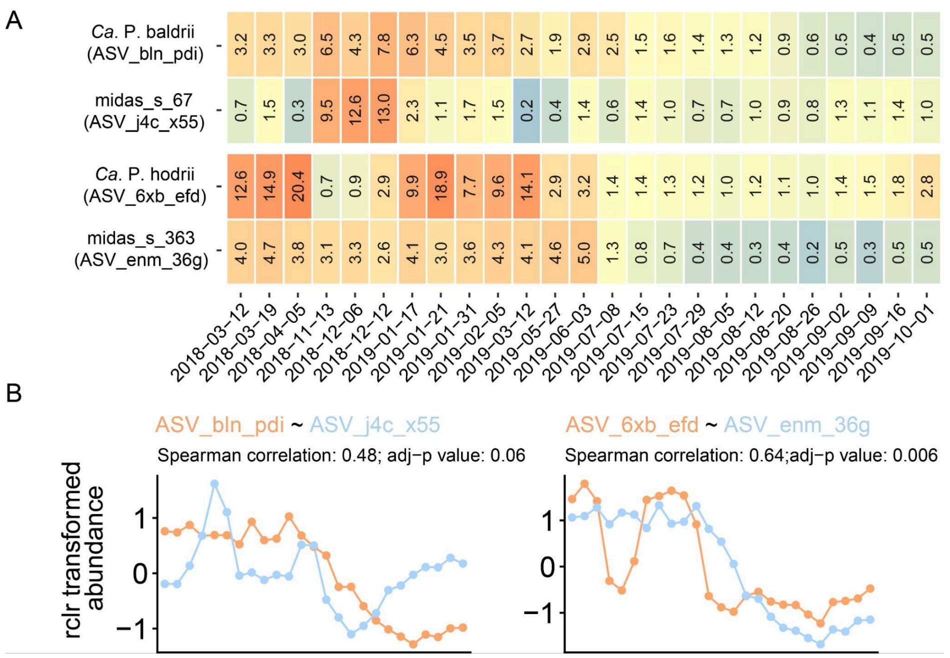


**Supplementary Fig. 5. Co-occurrence analyses of *Ca.* Lutisaccharimonas and *Ca.* Phosphoribacter in the Fredericia WWTP.** (A) Relative abundance of ASVs of *Ca*. P. baldrii, *Ca*. P. hodrii, *Ca*. Lutisaccharimonas ASV_j4c_x55 and *Ca*. Lutisaccharimonas

ASV_enm_36g during the sampling period. (B) Correlation plot of the rclr-transformed abundance of the ASV pairs of *Ca*. P. baldrii (ASV_bln_pdi) and *Ca.* Lutisaccharimonas ASV_j4c_x55 (left), *Ca*. P. hodrii and *Ca.* Lutisaccharimonas ASV_enm_36g (right) across the sampling period. The x-axis (sampling date) is shown in panel A.

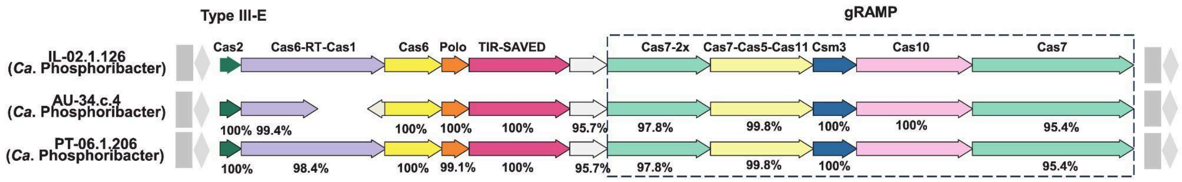


**Supplementary Fig. 6**. Genomic arrangement of the type III-E CRISPR locus in the three *Ca.* Phosphoribacter MAGs that were found to contain spacer sequences from *Ca*. Lutisaccharimonas. Different Cas proteins are shown in different colors. Cas protein sequence identities against IL-02.1.126 are shown as percentages below the arrows. IL–Italy; AU–Australia; PT–Portugal.

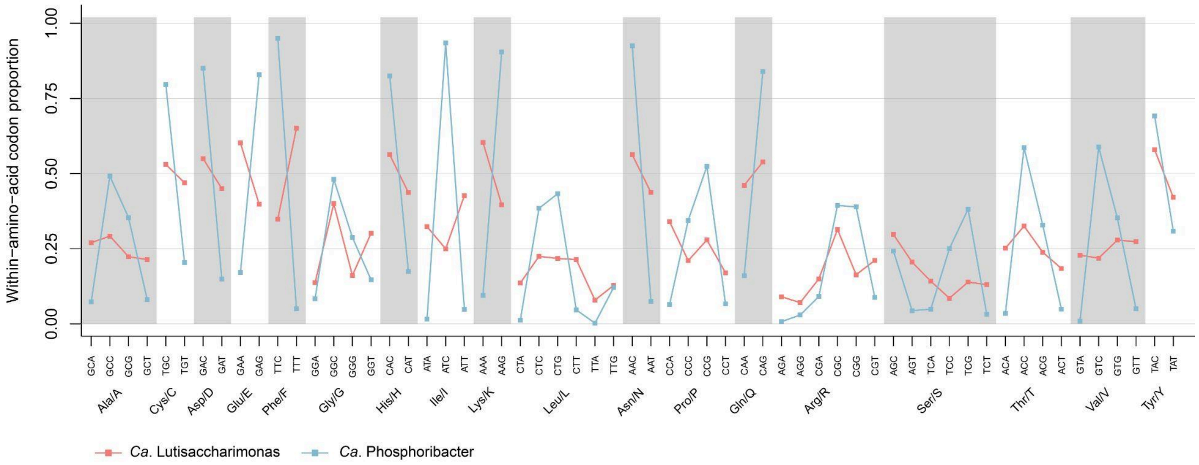


**Supplementary Fig. 7. Codon usage pattern in *Ca.* Lutisaccharimonas (GCA_016700375.1) and *Ca.* Phosphoribacter (GCA_016704565.1).** The y-value indicates the frequency of each synonymous codon for the respective amino acid.

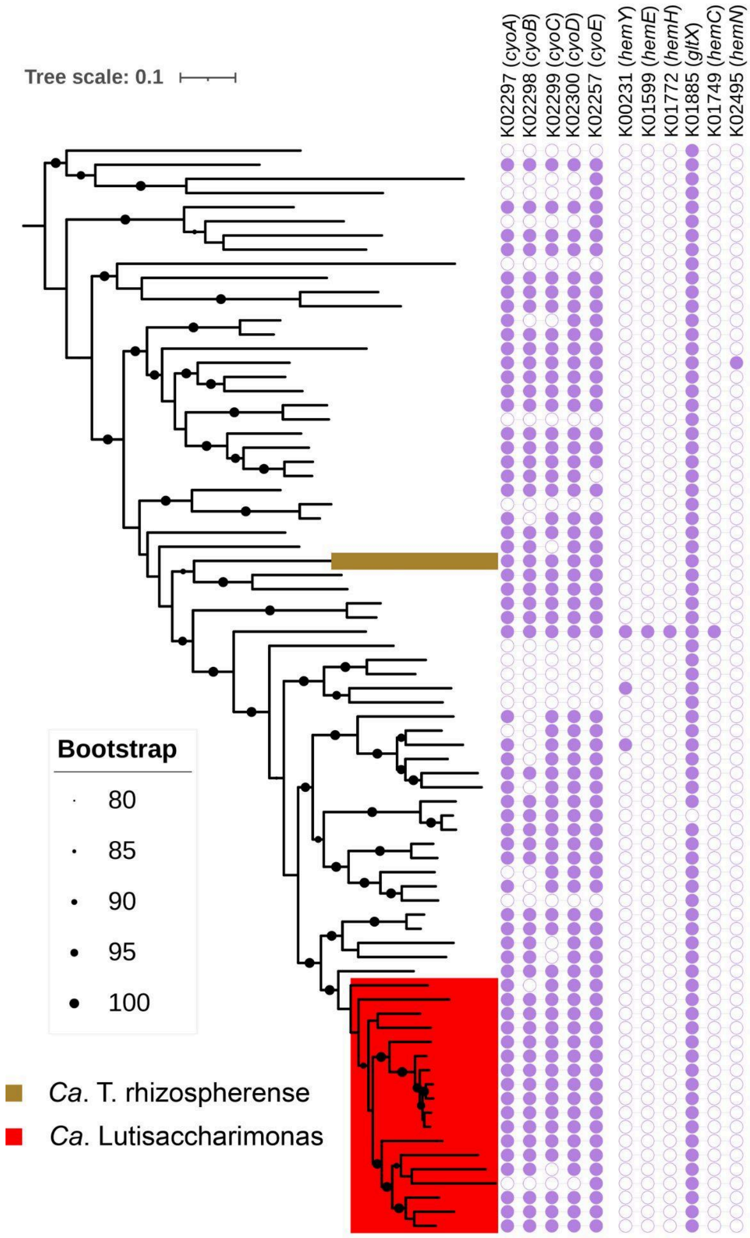


**Supplementary Fig. 8. Genes encoding cytochrome bo$_3$-type ubiquinol oxidase subunits I-IV (*cyoABCD*) and heme biosynthesis gene (*cyoE*) among family UBA4665 members within the *Saccharimonadia*.** The phylogenetic tree was constructed based on GTDB r220 single copy marker genes with IQ-TREE 2. *Ca*. Lutisaccharimonas MAGs within the UBA4665 family are highlighted in red, whereas *Ca*. Teamsevenus rhizospherense is labeled in brown.

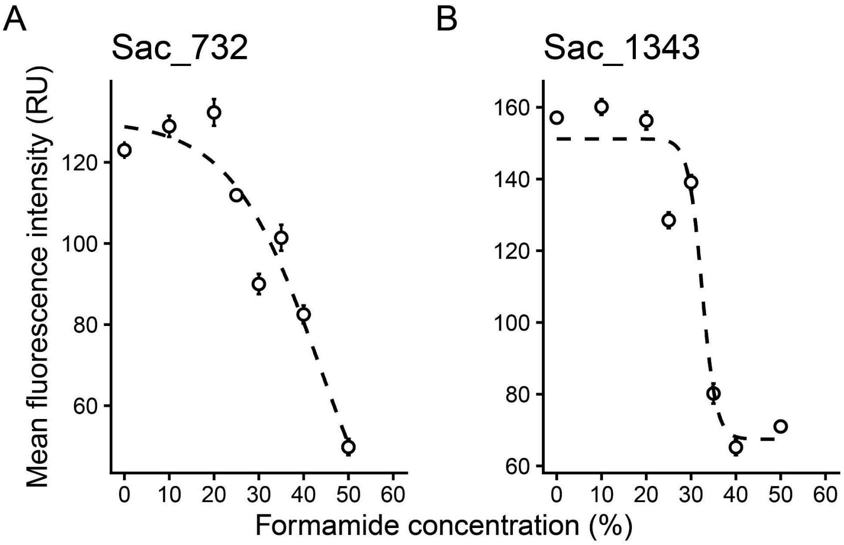


**Supplementary Fig. 9. Dissociation profiles of the newly designed 16S rRNA-targeted oligonucleotide probes (panel A) Sac732 and (panel B) Sac1343 targeting *Ca.* Lutisaccharimonas**. Per-cell probe fluorescence was measured after CloneFISH of *E. coli* cells. Data points depict mean per-cell intensity (error bars, s.e.m.). Stippled lines are the probe dissociation curves estimated by sigmoidal regression. Five randomly picked microscopic fields of view were recorded for each formamide concentration. RU, relative units.

### References


1. Özcan, A. *et al.* Programmable RNA targeting with the single-protein CRISPR effector Cas7-11. *Nature* **597**, 720–725 (2021).
2. Yu, G. *et al.* Structure and function of a bacterial type III-E CRISPR–Cas7-11 complex. *Nat Microbiol* **7**, 2078–2088 (2022).
3. Wang, J. Y. *et al.* Structural coordination between active sites of a CRISPR reverse transcriptase-integrase complex. *Nat Commun* **12**, 2571 (2021).
4. Hogrel, G. *et al.* Cyclic nucleotide-induced helical structure activates a TIR immune effector. *Nature* **608**, 808–812 (2022).
5. Shmakov, S. A. *et al.* CRISPR Arrays Away from cas Genes. *CRISPR J* **3**, 535–549 (2020).
6. Burstein, D. *et al.* Major bacterial lineages are essentially devoid of CRISPR-Cas viral defence systems. *Nat Commun* **7**, 10613 (2016).
7. Chen, L.-X. *et al.* Candidate Phyla Radiation Roizmanbacteria From Hot Springs Have Novel and Unexpectedly Abundant CRISPR-Cas Systems. *Front. Microbiol.* **10**, (2019).
8. Wang, J., Zhong, H., Chen, Q. & Ni, J. Adaption mechanism and ecological role of CPR bacteria in brackish-saline groundwater. *npj Biofilms Microbiomes* **10**, 141 (2024).
9. Castelle, C. J. *et al.* Biosynthetic capacity, metabolic variety and unusual biology in the CPR and DPANN radiations. *Nat Rev Microbiol* **16**, 629–645 (2018).
10. He, Y. *et al.* Candidate Phyla Radiation (CPR) bacteria from hyperalkaline ecosystems provide novel insight into their symbiotic lifestyle and ecological implications. *Microbiome* **13**, 94 (2025).

11. Starr, E. P. *et al.* Stable isotope informed genome-resolved metagenomics reveals that Saccharibacteria utilize microbially-processed plant-derived carbon. *Microbiome* **6**, 122 (2018).
12. Nicolas, A. M. *et al.* Soil Candidate Phyla Radiation Bacteria Encode Components of Aerobic Metabolism and Co-occur with Nanoarchaea in the Rare Biosphere of Rhizosphere Grassland Communities. *mSystems* **6**, 10.1128/msystems.01205-20 (2021).
13. Nierychlo, M. *et al.* MiDAS 3: an ecosystem-specific reference database, taxonomy and knowledge platform for activated sludge and anaerobic digesters reveals species-level microbiome composition of activated sludge. *Water Research* **182**, 115955 (2020).
